# Differential proteomic and metabolic remodelling of CD4⁺ and CD8⁺ T cells in mesenteric lymph nodes during colitis-associated colorectal cancer

**DOI:** 10.64898/2025.12.26.696593

**Authors:** Diego González-Romero, Laura Spinelli, Pilar Fajardo, Ester Díaz-Mora, Marta Gómez de Cedrón, Juan José Sanz-Ezquerro, Doreen A. Cantrell, Ana Cuenda

## Abstract

Patients with colitis are at elevated risk of developing colitis-associated colorectal cancer (CAC), yet the molecular and metabolic adaptations of T cells during this process remain unclear. Using quantitative mass spectrometry in the AOM/DSS mouse model we mapped proteomic networks of CD4^+^ and CD8^+^ T cells in mesenteric lymph nodes across healthy, colitis, and cancer stages. CD4⁺ T cells expanded during CAC progression and underwent extensive proteomic remodelling, including upregulation of DNA replication, cell cycle, ribosome biogenesis, and translation. These changes were accompanied by enhanced glycolysis, TCA activity, mitochondrial mass, and oxidative phosphorylation, indicating a highly metabolically active state. In contrast, CD8⁺ T cells showed minimal changes in cell number, protein content and in metabolic and proteomic remodelling. CD4⁺ T cells progressively acquire inflammatory Th1 and regulatory features, whereas CD8⁺ T cells remained poorly activated. Together, these findings reveal distinct T cell adaptations shaped by CAC and identify potential immunometabolic targets.

## Introduction

Colorectal cancer (CRC) is the second leading cause of cancer-related mortality worldwide, and its incidence has been rising dramatically among individuals under 50 years of age, making early-onset CRC an emerging global health concern^1,2^. Patients with colitis, inflammatory bowel disease (IBD), including ulcerative colitis and Crohn’s disease, are at increased risk of developing colitis-associated colorectal cancer (CAC)^3^. The pathogenesis of CAC involves a complex interplay between persistent inflammation, immune dysregulation, and epithelial transformation, driving malignant progression. Among the immune components implicated in this process, T cells play a central role in orchestrating both protective and pathological responses. CD4⁺ and CD8⁺ T cells are the most abundant immune cells in CRC tumours, and their activation, differentiation, and metabolic adaptation are critical for shaping the inflammatory milieu and influencing cancer progression^4,5^. Recent studies have highlighted the importance of T cell subsets in tumour immunity, particularly in CRC, where their functional diversity and spatial distribution within the tumour microenvironment appear to be key determinants of the clinical outcome^5^. The tumour microenvironment harbours subsets of T cells with divergent roles: anti-tumorigenic populations, such as effector CD8⁺ T cells or CD4⁺ Th1 cells, and pro-tumorigenic subsets, including CD4⁺ regulatory T cells (Tregs). Even CD8⁺ T cells exhibit heterogeneity, as they may undergo exhaustion, enter hyporesponsive states, or become dysfunctional^5^.

Despite extensive research on the immunological mechanisms underlying CAC, the specific molecular and metabolic changes occurring in T cells during the transition from colitis to colorectal cancer remain largely unknown. Understanding these changes, in the context of immunologically cold tumours like CRC^5^, is crucial for identifying therapeutic targets that can reprogram the immune microenvironment, enhance anti-tumour immunity, and convert these immunologically silent tumours into responsive ones. Most studies have focused on tumour-infiltrating lymphocytes, through phenotypic characterization using flow cytometry to define T cell subsets^5–7^. Transcriptomic profiling has also provided insights into gene expression patterns within the tumour microenvironment. However, a critical gap remains in our understanding of how CD4⁺ and CD8⁺ T cells are modulated in lymphoid organs such as the mesenteric lymph nodes (mLN), key sites for immune cell priming and activation in the gut^8^. During early tumour initiation, naïve T cells traffic to tumour-draining lymph nodes, where they are primed by activated antigen-presenting cells (APCs), such as dendritic cells (DCs). Antigen-specific stimulation of CD8⁺ cytotoxic and CD4⁺ helper T cells via the T cell receptor (TCR) initiates immune responses, driving the generation of effector populations capable of targeting cancer cells^9^. This is a critical step, since can shift the balance toward either anti-tumour effector T cells, or pro-tumour Treg cells^10^. Thus the immune status of mLN critically contribute to shape the tumour microenvironment, and tumours may suppress mLN function to evade immune responses^11^. To date, no comprehensive protein expression analysis has characterized the dynamics and proteomic landscape of CD4⁺ and CD8⁺ T cell populations in mLN during the transition from healthy to colitis, and to cancer.

In this study, we address this gap by providing a detailed proteomic characterization of CD4⁺ and CD8⁺ T cells in mLN during CAC progression. Using quantitative high-resolution mass spectrometry, we found distinct immunometabolic trajectories in these T cell subsets. Whereas CD4⁺ T cells undergo expansion and metabolic reprogramming, acquiring features of both inflammatory and regulatory phenotypes, CD8⁺ T cells exhibit limited proteomic changes, which may contribute to disease progression. These findings offer new insights into T cell adaptation in the context of chronic inflammation and tumorigenesis, highlighting potential targets for therapeutic intervention.

## Results

### Proteome remodelling of CD4⁺ and CD8⁺ T cells in mLN throughout CAC

As mesenteric lymph nodes (mLN) drain the tumour-bearing colon and play a central role in coordinating immune responses in the gut^8,12^, we selected this tissue to investigate proteomic changes in CD4⁺ and CD8⁺ T cells during colitis-associated cancer (CAC) progression. Specifically, we used the AOM/DSS mouse model, representative of inflammation-driven colorectal carcinogenesis and mimicking human sporadic CRC and CAC at histological and molecular levels^13^. This allowed the analysis of T cells at two key time points: the early stage of acute inflammation (colitis; day 15) and the late stage when mice have developed tumours (cancer; day 60) (Suppl. Fig. 1A-B). Consistent with previous reports describing mLN enlargement during colitis and CAC^14,15^, we found a significant increase in the total number of T cells and other immune cells in the mLNs of mice with colitis and CAC compared to healthy controls (Fig. 1A, 1B, Suppl. Fig. 1C-D). Of note, the absolute number of CD4⁺ T cells were significantly elevated in CAC, whereas CD8⁺ T cells remained comparable to control levels (Fig. 1B). These findings reflect a preferential expansion and/or recruitment of CD4⁺ T cells during inflammation and tumour development.

**Figure 1.**
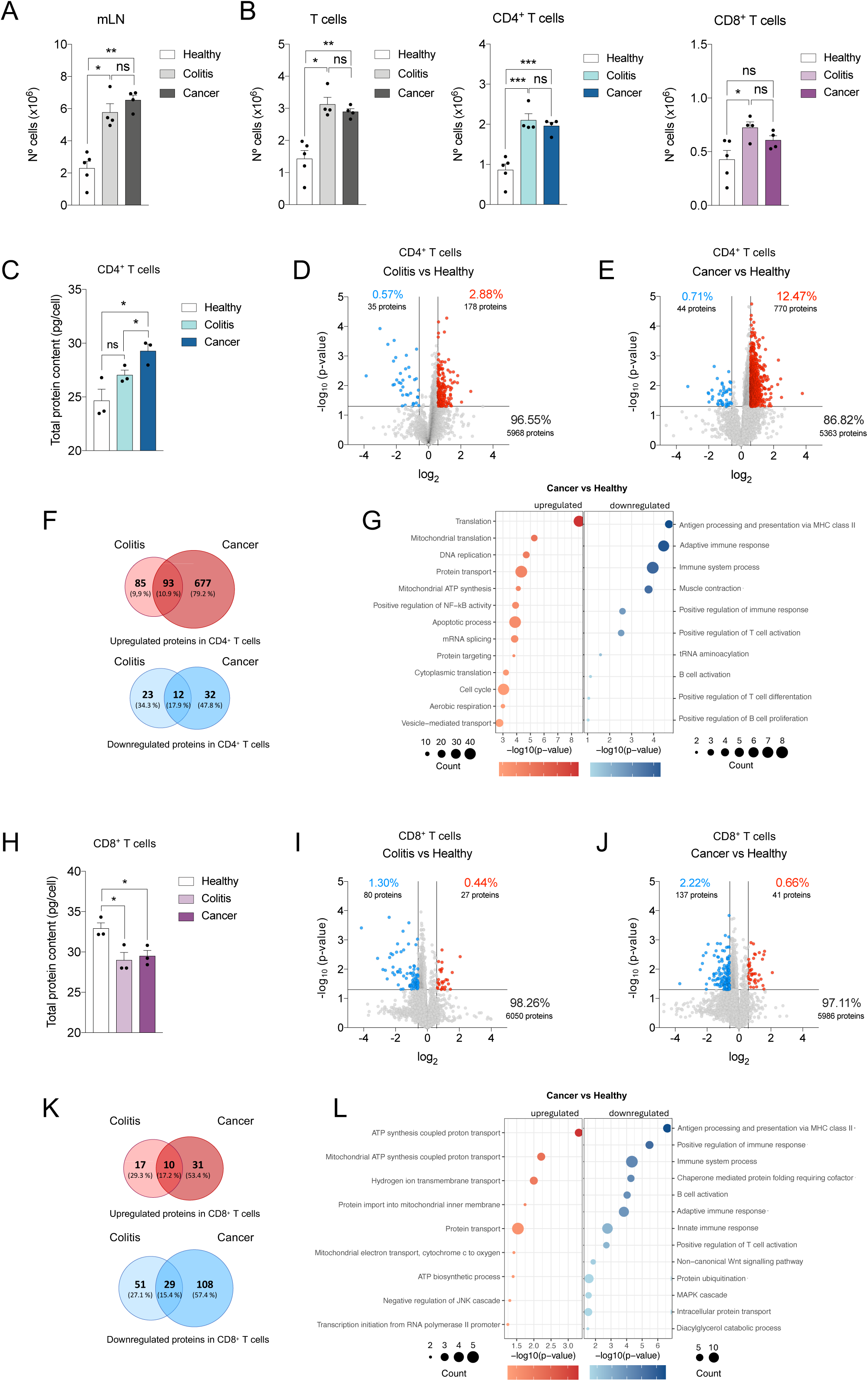
Proteome remodelling of CD4^+^ and CD8^+^ T cells in mLN throughout CAC. Mice were treated with AOM/DSS, and at days 0 (Healthy), 15 (Colitis) and 60 (Cancer), CD4^+^ and CD8^+^ T cells were sorted from mLN and their proteomes analysed by quantitative mass spectrometry. **(A)** Total mLN cell number, **(B)** Total T cell number (left), CD4^+^ (centre) and CD8^+^ T cells (right) in the mLN of healthy, colitis and CAC mice. Each dot represents a single mouse (n=4-5). **(C)** Total protein content in CD4^+^ T cells. Each dot represents a pool of three mice (n=3 pools). **(D-E)** Volcano plots showing the difference in the expression of proteins between (D) Colitis and Healthy CD4^+^ T cells and (E) Cancer and Healthy CD4^+^ T cells. The horizontal lines indicate a *p*-value of 0.05. The vertical lines indicate a fold change of ± 1.5. The percentage and number of proteins that are downregulated (blue), upregulated (red), or unchanged (grey) are indicated. **(F)** Venn diagrams showing the overlaps of proteins upregulated or downregulated in CD4^+^ T cells from mLN over Colitis and Cancer relative to Healthy. **(G)** Bubble plot showing the enrichment analysis of GO biological processes of the differentially upregulated (left) or downregulated (right) proteins in Cancer versus Healthy in CD4^+^ T cells. Bubble size indicates the number of genes involved in each process, and bubble colour reflects the fold change in Cancer over Healthy conditions. **(H)** Total protein content in CD8^+^ T cells along CAC progression. Each dot represents a pool of three mice (n=3 pools). **(I-J)** Volcano plots showing the difference in the expression of proteins between (I) Colitis and Healthy CD8^+^ T cells and (J) Cancer and Healthy CD8^+^ T cells. The horizontal lines indicate a *p*-value of 0.05. The vertical lines indicate a fold change of ± 1.5. The percentage and number of proteins that are downregulated (blue), upregulated (red), or unchanged (grey) are indicated. **(K)** Venn diagrams showing the overlap of proteins upregulated or downregulated in CD8^+^ T cells Colitis and Cancer relative to Healthy. **(L)** Bubble plot showing the enrichment analysis of GO biological processes of the differentially upregulated (left) or downregulated (right) proteins in Cancer versus Healthy in CD8^+^ T cells. Bubble size indicates the number of genes involved in each process, and bubble colour reflects the fold change in Cancer versus Healthy conditions. All the data are shown as mean ± SEM. In all panels, ns = non-significant, \**p* ≤ 0.05, \*\**p* ≤ 0.01, \*\*\**p* ≤ 0.001.

Quantitative mass spectrometry analysis of the proteomes of CD4^+^ and CD8^+^ T cells isolated from mLN identified > 6000 proteins; protein mass and copy number per cell were estimated by “proteomic ruler” method^16^. Our data revealed a clear increase in total protein content in CD4^+^ T cells during disease progression, which was particularly marked in cancer, with approximately a 12.5% change in protein expression compared to healthy controls (Fig. 1C-F). Proteins that were significantly upregulated in these cells in both cancer and colitis were primarily involved in translation, DNA replication, cell cycle and mitochondrial ATP synthesis (Fig. 1G, Suppl. Fig. 2A). These findings indicate elevated CD4^+^ T cell activation and metabolic demand.

In contrast, CD8^+^ T cells exhibited a decrease in total protein content following AOM/DSS treatment (Fig. 1H). Interestingly, their proteome remained relatively stable throughout disease progression, with up to 98% similarity across conditions (Fig. 1I-K). Since CD8^+^ T cells typically play a crucial role in anti-tumour immunity in CAC, the lack of significant proteomic changes, along with reduced expression of key proteins involved in T and B cell activation and innate immune responses (Fig. 1L, Suppl. Fig. 2B), may reflect functional impairment and weakened coordination with other immune cells, contributing to immune evasion in the tumour microenvironment. This also indicate a limited adaptive immune response against developing tumours.

A significant finding with potential implications for CAC was the consistent downregulation of proteins involved in antigen processing and presentation via MHC-II in both CD8^+^ and CD4^+^ T cells (Fig. 1G, 1L, Suppl. Fig. 2A 2B), which reaches its maximum in colitis and remains low in cancer (Suppl. Fig. 3A-B). This observation might reflect a loss of antigen-presenting capacity, contributing to an immunosuppressive microenvironment that favours tumour progression from early stages of CAC. This effect was corroborated by flow cytometry analysis of CD4^+^ and CD8^+^ T cells in colitis, showing a marked reduction in the percentage of positive cells and a significant decrease in the median fluorescence intensity (MFI), indicating lower expression levels per cell (Suppl. Fig. 3C-F).

### Upregulation of ribosomal and initiation complex proteins in CD4⁺ T cells in CAC

The increase in protein content observed in CD4⁺ T cells during CAC implies an enhancement in protein synthesis, correlating with a notable increase in ribosome-related protein mass over time, particularly during the first 15 days of AOM/DSS treatment (Fig. 2A). Ribosomes are macromolecular machines, composed by ribosomal RNA and many ribosomal proteins, that facilitate protein synthesis. Accordingly, we found a general upregulation of the amount of ribosomal proteins and of proteins involved in the protein synthesis machinery in CD4⁺ T cells from mice having colitis and cancer (Fig. 2B, 2C). Accordingly, the expression of amino acid transporters was induced during CAC progression (Fig. 2D), supporting amino acid intake to meet demands of enhanced protein synthesis. Expression of key amino acid transporters such as SLC7A5/SLC3A2 (Met, Leu, Iso, Trp and Val) and SLC38A1 (Gln and Met) increased progressively over CAC development (Fig. 2E).

**Figure 2.**
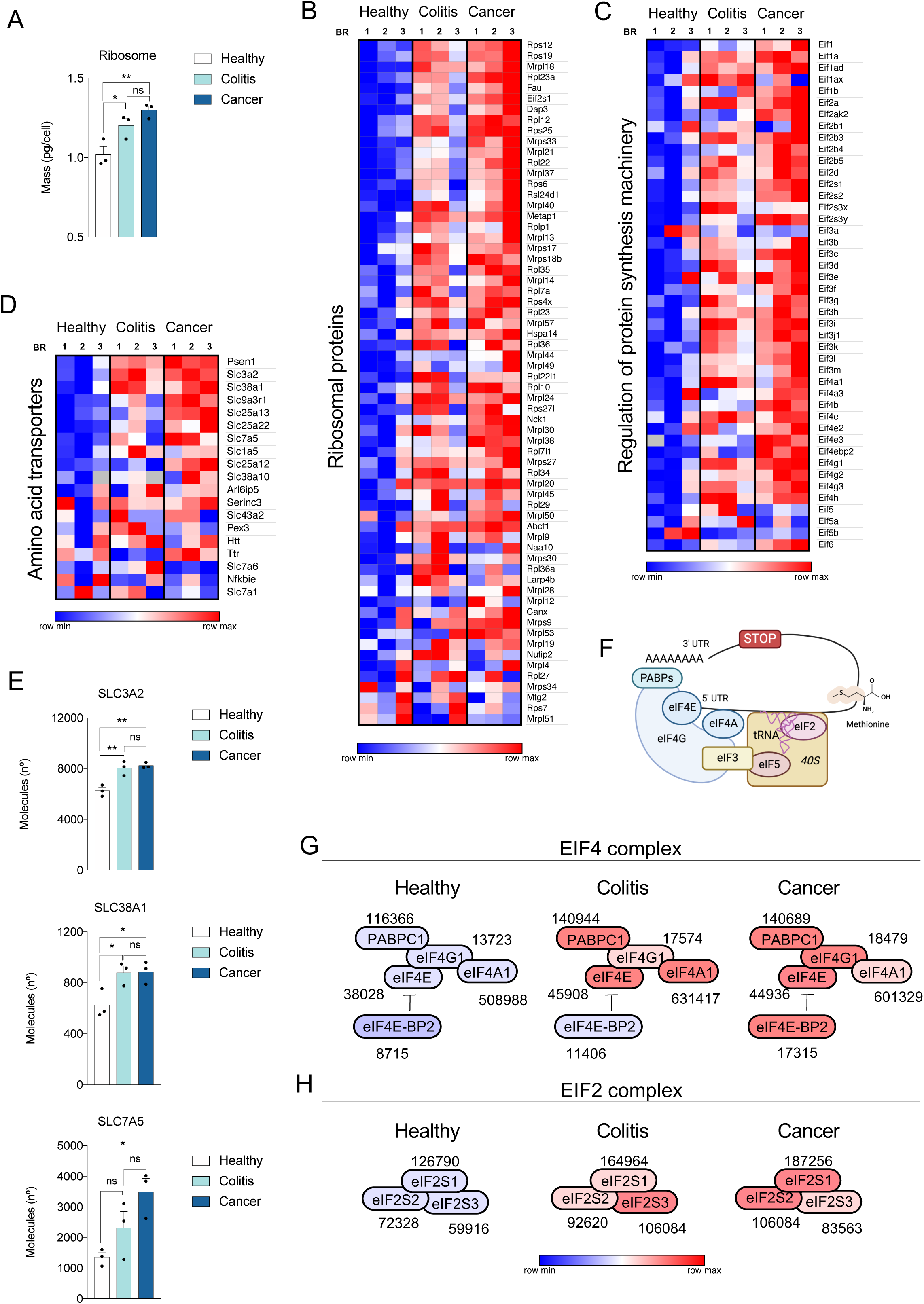
Upregulation of ribosomal and initiation complex proteins in CD4^+^ T cells in CAC. Mice were treated with AOM/DSS, and at days 0 (Healthy), 15 (Colitis) and 60 (Cancer), CD4^+^ T cells were sorted from mLN and their proteomes analysed by quantitative mass spectrometry. **(A)** Total ribosome mass in CD4^+^ T cells. Each dot represents a pool of three mice (n=3 pools). **(B-D)** Heat maps of proteins involved in (B) ribosomal proteins (GO:0005840), (C) regulation of protein synthesis machinery and (D) amino acid transporters (GO:0006865) identified in CD4^+^ T cells proteomes during CAC progression. BR, biological replicate. Relative protein abundance is graded from low (blue) to high (red) to allow comparisons between different CAC stages. **(E)** Protein copy numbers per cell (Molecules (n°)) of the amino acid transporters SLC3A2, SLC38A1 and SL7A5 in CD4^+^ T cells. Each dot represents a pool of three mice (n=3 pools). **(F)** Schematic representation of translation initiation complex. Image created with BioRender. **(G-H)** Stoichiometric models for (G) EIF4 and (H) EIF2 complexes. (G) Protein copy numbers for PABPC1, eIF4G1, eIF4E, eIF4A1 and eIF4E-BP2 in CD4^+^ T cells are shown as numbers. (H) Protein copy numbers for eIF2S1, eIF2S2 and eIF2S3 in CD4^+^ T cells are shown as numbers. The data represented is the average value of three pooled samples, each consisting of three mice (n=3 pools; 9 mice in total). Relative protein abundance is graded from low (blue) to high (red). All the data are shown as mean ± SEM. In all panels, ns = non-significant, \**p* ≤ 0.05, \*\**p* ≤ 0.01.

A major point of the regulation of protein synthesis is at the translation initiation. In mammalian cells, mRNAs are recruited to the ribosomes via the eukaryotic initiation factors 4F (eIF4F) complex, which comprises the proteins eIF4E, eIF4G and eIF4A. This complex delivers mRNA to the preinitiation complex, which includes eIF2 and mediates the binding of initiator tRNA^17,18^ (Fig. 2F). Our analyses reveal a marked increase in the abundance of proteins from these complexes in colitis, being more pronounced in cancer (Fig. 2G, 2H). The upregulation of the EIF2 and EIF4 complexes seems to underlie the enhanced translation capacity in CD4^+^ T cells. This shift in protein synthesis machinery reflects broader changes in cellular function and correlates with the progressive rise in total protein content as CD4^+^ T cells transition from healthy states to inflammatory conditions and ultimately to malignancy.

### Changes in proteins involved in DNA replication and cell cycle during CAC

The observed increase in CD4^+^ T cell numbers during colitis and cancer indicates enhanced proliferation, correlating with the upregulated expression of DNA replication and cell cycle-related proteins detected in the proteomic analysis. A closer examination of these proteins revealed that those involved in DNA replication, such as MCM helicases, DNA polymerase complexes (POLA, POLD, POLE), replication protein A (RPA), replication factors 2 and 3 (RFC2 and RFC3), or ligase 1 (LIG1), were consistently upregulated during the colitis phase (first 15 days of treatment), with their levels remaining elevated in cancer (Fig. 3A).

**Figure 3.**
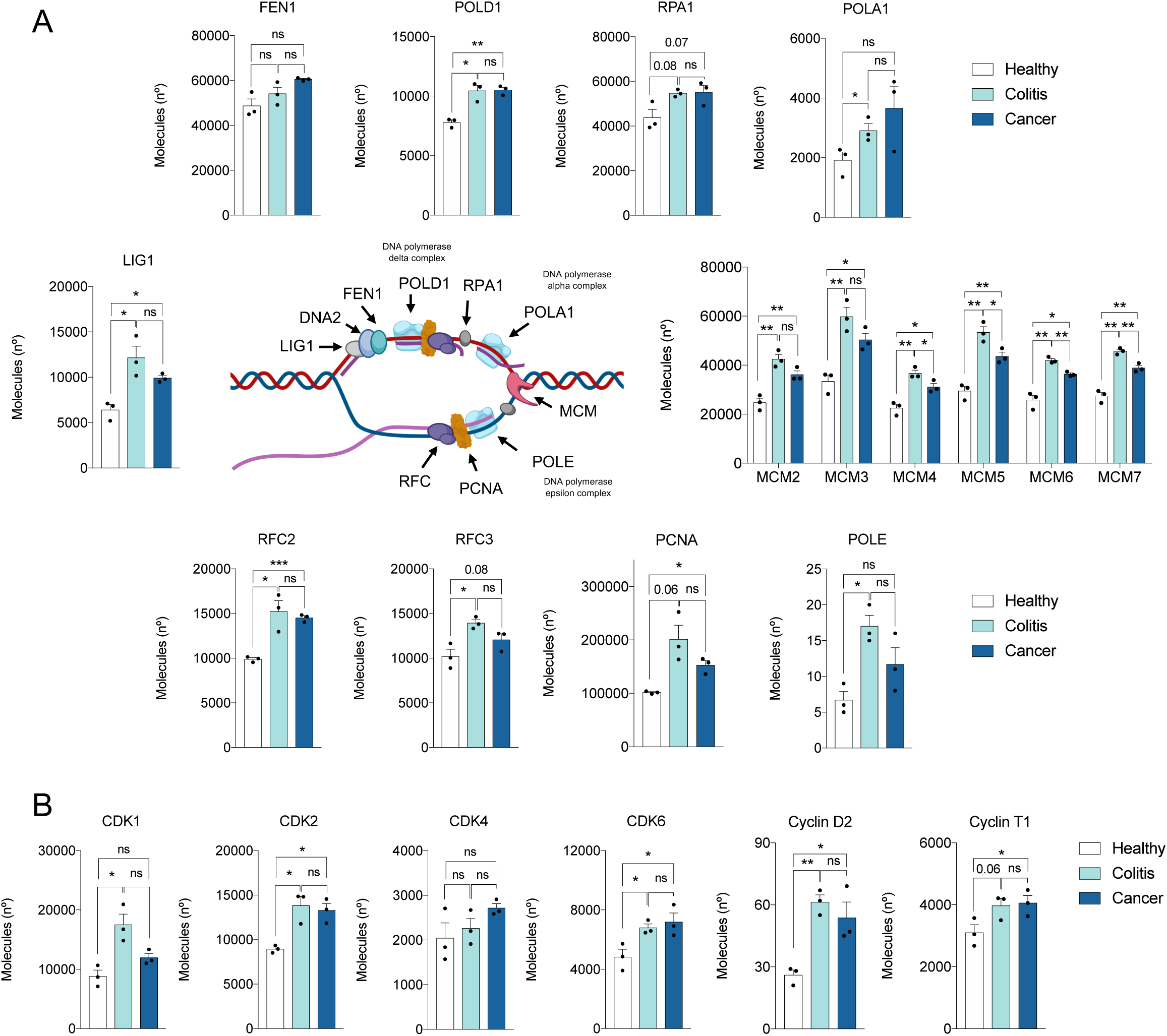
Changes in proteins involved in DNA replication and cell cycle during CAC. CD4^+^ T cells were sorted from mLN of AOM/DSS-treated mice at days 0 (Healthy), 15 (Colitis) and 60 (Cancer) and their proteomes analysed by quantitative mass spectrometry. **(A-B)** Expression profiles of key components involved in (A) DNA replication and (B) cell-cycle and transcriptional processes during CAC stages. Data are represented as protein copy numbers per cell (Molecules (n°)) in CD4^+^ T cells. Each dot represents a pool of three mice (n=3 pools). All the data are shown as mean ± SEM. In the centre of panel (A) is a schematic image of DNA replication fork and its associated processing machinery (generated with Biorender). In all panels, ns = non-significant, \**p* ≤ 0.05, \*\**p* ≤ 0.01, \*\*\**p* ≤ 0.001.

Furthermore, cyclin-dependent kinases (CDK1/2, CDK4/6) and cyclins (D and T), which regulate cell cycle progression and DNA transcription, also showed marked upregulation during colitis, with levels remaining higher in cancer compared to healthy conditions (Fig. 3B). These molecular changes support CD4^+^ T cell proliferation, observed as early as the colitis stage.

In contrast to CD4^+^ T cells, CD8^+^ T cells showed minimal changes of the proteins involved in proliferation and cell cycle regulation throughout the progression of CAC (Suppl. Fig. 4). Apart from an increase in cyclin D and a transient decrease in RFC2 and CDK1, most factors remained stable, consistent with the constant CD8^+^ T cell numbers observed in mLN throughout CAC development (Fig. 1B). This differential regulation suggests that early CD4^+^ T cell expansion may drive inflammation and tumour development, while CD8^+^ T cell stability could reflect a weakened anti-tumour response.

### Regulation of glucose metabolism and aerobic capacity of T cells during CAC

Protein synthesis, cellular proliferation, and cell signalling during T cell activation require substantial energy and building blocks produced through different metabolic pathways. Activated T cells increase glycolysis and oxidative phosphorylation (OxPhos) to generate ATP and metabolites that sustain proliferation while maintaining redox balance. Additionally, activated T cells rely mainly on glutaminolysis to fuel OxPhos, while decreasing mitochondrial fatty acid oxidation^19,20^. Our data showed that glycolysis-related enzymes in CD4^+^ T cells were markedly higher in colitis and cancer compared to healthy conditions (Fig. 4A-D), whereas glucose transporters (SLC2A1 and SLC2A3) and lactate transporter (SLC16A1) remained stable (Fig. 4E, 4F). Of relevance, the key amino acid transporter SLC38A1, upregulated during CAC (Fig 2E) to sustain protein synthesis in CD4⁺ T cell, also facilitates glutamine uptake as an anaplerotic substrate for the TCA cycle, supporting its role in meeting the high metabolic demands of CD4⁺ T cell proliferation.

**Figure 4.**
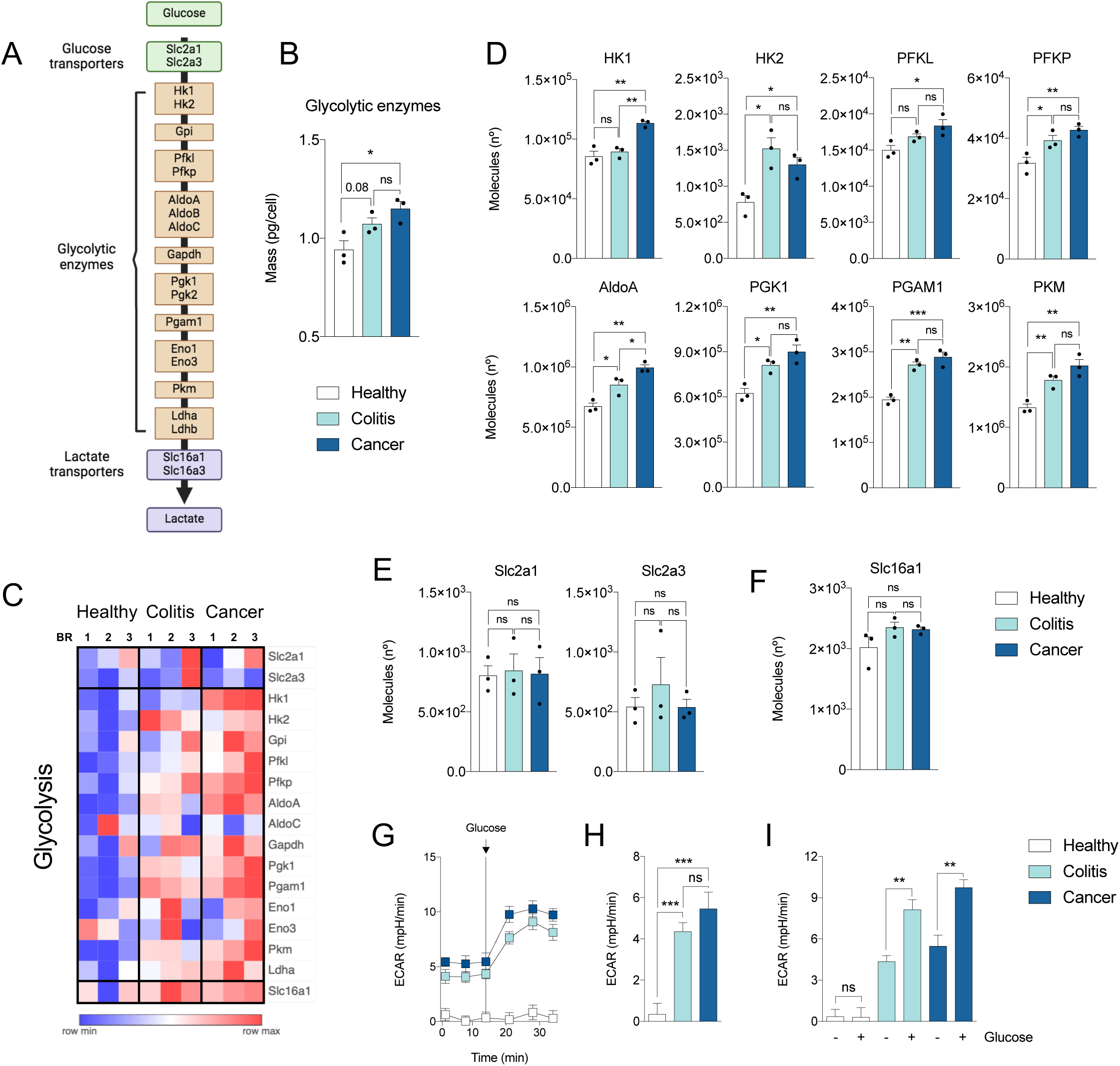
Regulation of glucose metabolism of T cells during CAC. Mice were treated with AOM/DSS, and at days 0 (Healthy), 15 (Colitis) and 60 (Cancer), CD4^+^ T cells were sorted from mLN and their proteomes analysed by quantitative mass spectrometry. **(A)** Schematic representation of anaerobic glycolysis. Image created with BioRender. **(B)** Total glycolytic enzymes mass per cell in CD4^+^ T cells. Each dot represents a pool of three mice (n=3 pools). **(C)** Heat map of proteins involved in glycolysis (GO:0006096) identified in CD4^+^ T cells proteomes in healthy, colitis, and cancer mice. BR, biological replicate. Relative protein abundance is graded from low (blue) to high (red) to allow comparisons. **(D)** Expression profiles of glycolytic enzymes during CAC progression. Data are represented as protein copy numbers per cell (Molecules (n°)) in CD4^+^ T cells. Each dot represents a pool of three mice (n=3 pools). **(E-F)** Abundances of (E) the glucose transporters SCL2A1 and SLC2A3 and (F) the lactate transporter SLC16A1 in CD4^+^ T cells. Data are represented as protein copy numbers per cell (Molecules (n°)). Each dot represents a pool of three mice (n=3 pools). **(G)** Real-time monitoring of extracellular acidification rate (ECAR) in CD4^+^ T cells along CAC progression. Arrow indicates the time of injection of glucose. **(H)** ECAR profile of CD4^+^ T cells in basal state in healthy, colitis, and CAC (n= 4-5 mice). **(I)** ECAR profile of CD4^+^ T cells before (-) and after (+) glucose treatment (n= 4-5 mice per condition). All the data are shown as mean ± SEM. In all panels, ns = non-significant, \**p* ≤ 0.05, \*\**p* ≤ 0.01, \*\*\**p* ≤ 0.001.

The metabolic reprogramming observed in CD4⁺ T cells during CAC was validated through functional studies, including measurements of extracellular acidification rate (ECAR) and oxygen consumption rate (OCR), indirect indicators of aerobic glycolysis and OxPhos. ECAR of CD4^+^ T cells from colitis and cancer showed higher basal glycolysis and strong responsiveness to glucose compared to CD4^+^ T cells from healthy mLN (Fig. 4G-I). Healthy mLN-CD4^+^ T cells not only exhibited lower ECAR rates but also failed to respond to glucose stimulation (Fig. 4G-I). Collectively, these results indicate that CD4^+^ T cells undergo pronounced glycolysis during CAC, enhancing glucose utilization to meet the energy demands of proliferation and effector activity.

CD8^+^ T cells, on the other hand, showed a different pattern. Protein mass implicated in glycolysis remained relatively stable throughout the AOM/DSS treatment, with no significant differences among healthy, colitis, and cancer conditions (Suppl. Fig. 5A-D). ECAR measurements confirmed these findings. Glycolitic capacity of CD8⁺ T cells from colitis and cancer remained similar to basal levels, and only CD8⁺ T cells from healthy mLN showed a significant increase in glycolytic activity upon glucose stimulation (Suppl. Fig. 5E-G). These data suggest that CD8^+^ T cells in both colitis and cancer conditions have an impaired glycolytic capacity compared to their healthy counterparts, and suggest that, unlike CD4⁺ T cells, do not undergo substantial metabolic reprogramming during CAC.

We then examined the aerobic capacity of T cells across CAC and found significant changes in the expression of proteins involved in the tricarboxylic acid (TCA) cycle. CD4^+^ T cells showed a marked increase in the expression of citrate synthase (CS), isocitrate dehydrogenase (IDH1), alpha-ketoglutarate dehydrogenase (OGDH), succinyl CoA synthetase (SUCLA2), and succinate dehydrogenase (SDHA), at day 60 of the treatment compared to day 0 (healthy) and/or day 15 (colitis) (Fig. 5A, 5B). This upregulation correlated with increased mitochondrial mass of these cells (Fig. 5C) and a shift towards enhanced oxidative phosphorylation (OxPhos) as disease progresses to cancer (Fig. 5D, 5E), together with higher ATP synthase levels (Fig. 5F), indicating boosted mitochondrial ATP production and overall metabolic activity in CD4^+^ T cells.

**Figure 5.**
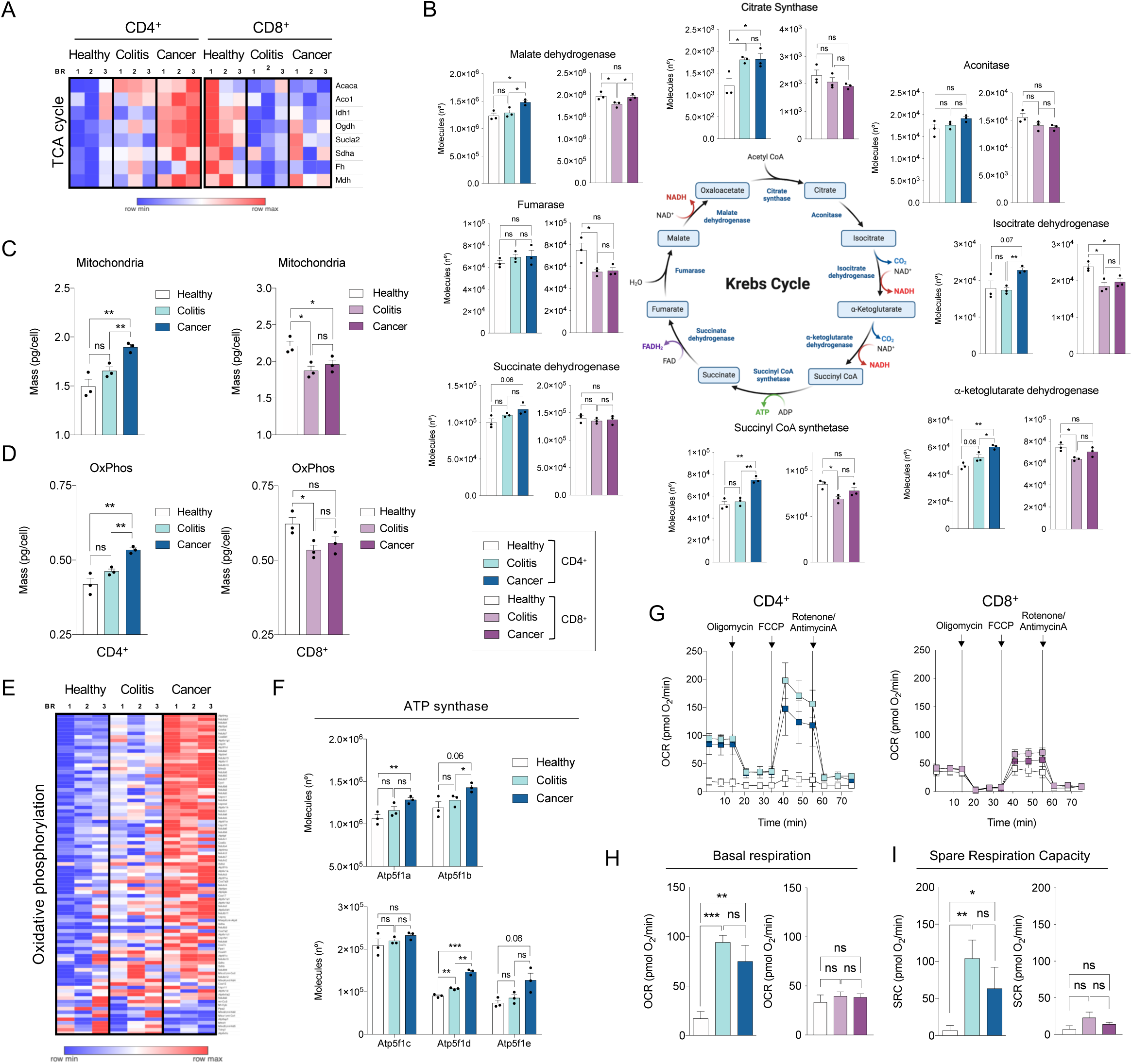
Regulation of aerobic capacity of T cells during CAC. CD4^+^ and CD8^+^ T cells were sorted from mLN at days 0 (Healthy), 15 (Colitis) and 60 (Cancer) from AOM/DSS-treated mice and their proteomes analysed by quantitative mass spectrometry. **(A)** Heat map of proteins involved in the tricarboxylic acid (TCA) cycle identified in CD4^+^ and CD8^+^ T cells proteomes during CAC progression. BR, biological replicate. Relative protein abundance is graded from low (blue) to high (red) to allow comparisons. **(B)** Expression profiles of key enzymes involved in the TCA/Krebs cycle in T cells in healthy, colitis, and cancer mice. Data are represented as protein copy numbers per cell (Molecules (n°)) in CD4^+^ and CD8^+^ T cells. Each dot represents a pool of three mice (n=3 pools). **(C-D)** (C) Total mitochondria mass and (D) Total OxPhos-related protein mass per cell in CD4^+^ (left) and CD8^+^ (right) T cells. Each dot represents a pool of three mice (n=3 pools). **(E)** Heat map of proteins involved in oxidative phosphorylation process (GO:0006119) identified in CD4^+^ T cells proteomes in healthy, colitis, and cancer. BR, biological replicate. Relative protein abundance is graded from low (blue) to high (red) to allow comparisons. **(F)** Protein copy numbers per cell (Molecules (n°)) of the ATP synthase proteins ATP5F1A, ATP5F1B, ATP5F1C, ATP5F1D and ATP5F1E in CD4^+^ T cells. Each dot represents a pool of three mice (n=3 pools). **(G)** Real-time monitorization of oxygen consumption rate (OCR) in CD4^+^ (left) and CD8^+^ (right) T cells (n= 3-5 mice per condition). Arrows indicate the time of injection of oligomycin, FCCP (Trifluoromethoxy carbonylcyanide phenylhydrazone) and rotenone/antimycin A. **(H)** OCR profile of CD4^+^ (left) and CD8^+^ (right) T cells in basal state in healthy, colitis, and CAC (n= 3-5 mice per condition). **(I)** Spare respiratory capacity (SRC) profile of CD4^+^ (left) and CD8^+^ (right) T cells in healthy, colitis and cancer (n= 3-5 mice per condition). All the data are shown as mean ± SEM. In all panels, ns = non-significant, \**p* ≤ 0.05, \*\**p* ≤ 0.01, \*\*\**p* ≤ 0.001.

In contrast, CD8^+^ T cells exhibited a different metabolic profile: several TCA cycle-related proteins were downregulated throughout CAC (Fig. 5A, 5B), and both mitochondrial mass and OxPhos were decreased at 15 and 60 days compared to healthy conditions (Fig. 5C, 5D), indicating metabolic impairment in these cytotoxic T cells as the disease advances.

To validate these findings on T cell metabolism at different stages of CAC, we analysed their bioenergetic profiles by measuring oxygen consumption rate (OCR), an OxPhos indicator, at a steady state and after inducing mitochondrial stress with modulators of respiration as described in Methods (Fig. 5G). CD4^+^ T cells from colitis and cancer animals, unlike CD8^+^ T cells, showed significantly higher basal respiration and spare respiratory capacity (SRC) compared to healthy (Fig 5H, 5I). This indicates greater metabolic flexibility and activity of CD4^+^ T cells in disease-associated conditions, likely supporting their activation and effector functions.

### Analysis of proteins involved in transcriptional and functional reprogramming of T cells in CAC

Proteome remodelling in T cell activation depends on mRNA synthesis, requiring activation and expression of transcription factors critical for T cell function. We analysed transcription factors and proteins implicated in T cell activation and found increased expression from healthy to cancer in CD4^+^ T cells (Fig. 6A), but not in CD8^+^ T cells (Suppl. Fig. 6A). This was further supported in CD4^+^ T cells by upregulation of key proteins downstream of the TCR, such as MALT1 and B-RAF, which likely enhance NFκB and MAPK signalling pathways, respectively, leading to activation of transcription factors like NFκB and AP-1 that drive T cell proliferation, cytokine production and effector functions (Fig. 6A, 6B)^21,22^. Accordingly, we also detected increased expression of transcription factors including NFκB, ATF2, CREB and NFATc (Fig. 6A, 6B). In contrast, SLA2, a negative regulator of TCR signalling^23^, was downregulated (Fig. 6A, 6B).

**Figure 6.**
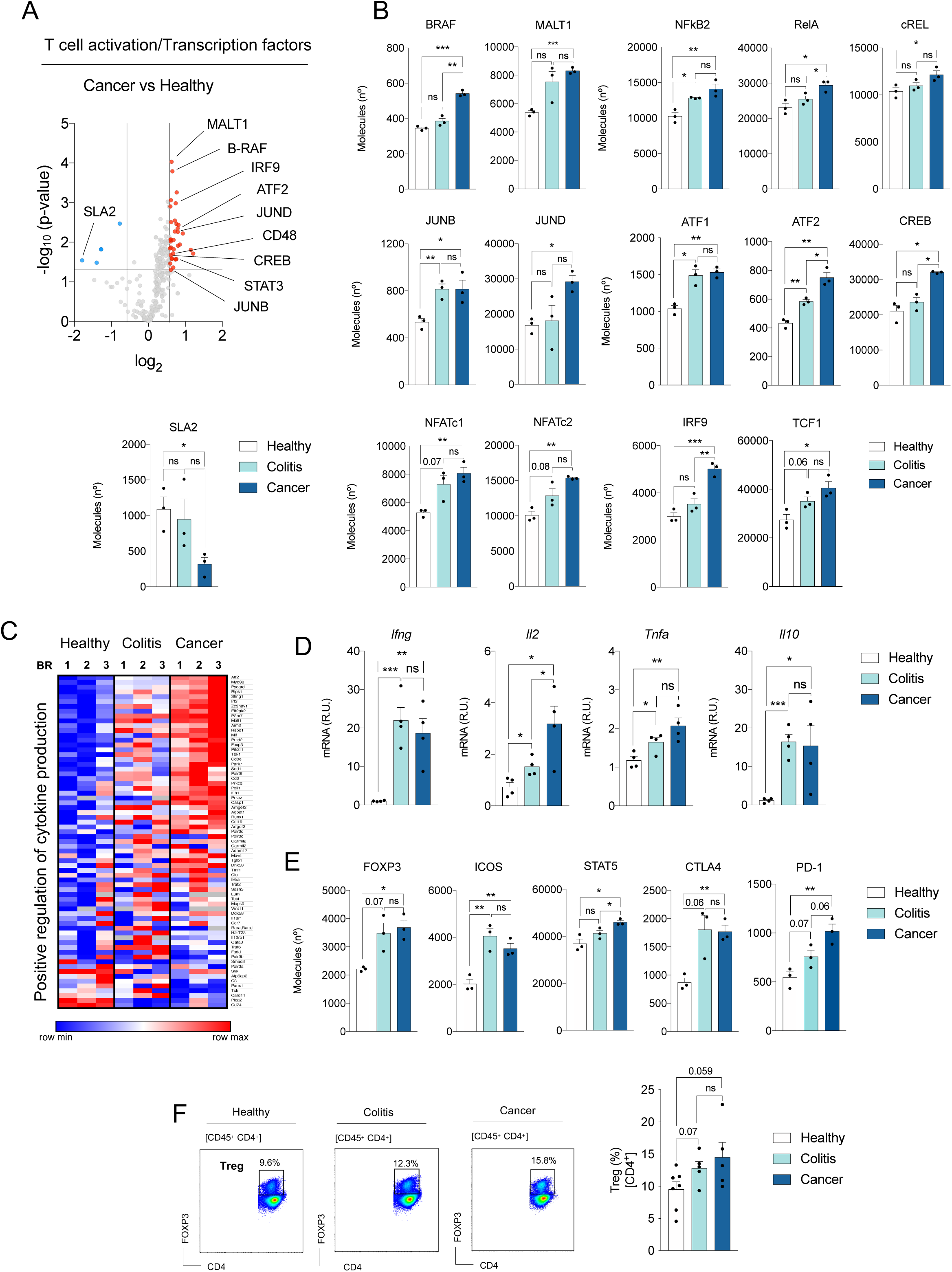
Analysis of proteins implicated in T cell activity during CAC progression. Mice were treated with AOM/DSS, and at days 0 (Healthy), 15 (Colitis) and 60 (Cancer), CD4^+^ T cells were sorted from mLN and their proteomes analysed by quantitative mass spectrometry. **(A)** Volcano plot showing the differential expression of proteins associated with T cell activation (GO:0042110) and DNA-binding transcription factor activity (GO:0003700) in Cancer versus Healthy CD4^+^ T cells. The horizontal lines indicate a *p*-value of 0.05. The vertical lines indicate a fold change of ± 1.5. **(B)** Expression profiles of transcription factors related to T cell activation during CAC progression. Data are represented as protein copy numbers per cell (Molecules (n°)) in CD4^+^ T cells. Each dot represents a pool of three mice (n=3 pools). **(C)** Heat map of proteins associated with the “positive regulation of cytokine production” (GO:0001819) identified in CD4^+^ T cells proteomes during CAC progression. BR, biological replicate. Relative protein abundance is graded from low (blue) to high (red) to allow comparisons. **(D)** Relative mRNA expression of the different genes at the indicated CAC stages in CD4^+^ T cells. *Actb* was used as control. Each dot represents a single mouse (n= 4). R.U.= relative units. **(E)** Copy numbers per cell (Molecules (n°)) of CD4^+^ Treg-associated markers FOXP3, ICOS, STAT5, CTLA4, and PD-1. Each dot represents a pool of three mice (n=3 pools). **(F)** Representative flow cytometry plots (Healthy, Colitis and Cancer) and percentages of regulatory CD4^+^ T cells (CD45^+^ CD4^+^ Foxp3^+^) in mLN from healthy, colitic and cancer-bearing mice. Each dot represents a single mouse (n=3-5). All the data are shown as mean ± SEM. In all panels, ns = non-significant, \**p* ≤ 0.05, \*\**p* ≤ 0.01, \*\*\**p* ≤ 0.001.

To further explore the functional implications of these transcriptional changes, we analysed proteins associated with the ’Positive regulation of cytokine production’ in CD4⁺ T cells. We observed a gradual increase in their expression from the healthy to cancer (Fig. 6C), which may contribute to the enhanced cytokine production capacity of these cells during cancer progression. Consistently, cytokine gene expression analysis showed that the levels of *Il2* and *Tnfa* mRNA produced by CD4⁺ T cells peak at day 60, whereas *Ifng* and *Il10* rise during colitis and remain elevated in cancer (Fig. 6D).

In parallel, we observed increased expression of Foxp3, ICOS and STAT5, markers of regulatory CD4⁺ T cell (Treg) in CAC (Fig. 6E), suggesting an enrichment of this immunosuppressive population in mLN. Flow cytometry analysis confirmed the increase in Foxp3⁺ regulatory T cells in mLN in mice with CAC (Fig. 6F). Moreover, enhanced levels of inhibitory receptors such as CTLA4 and PD-1 (Fig. 6E) reinforce their suppressive capacity, pointing to an active role of Treg in shaping an immunosuppressive microenvironment, limiting effector T cell responses and potentially facilitating tumour progression.

Interestingly, dendritic cells (DCs) showed one of the most pronounced increase in the mLN of mice with CAC (Suppl. Fig. 6B-C). Among conventional DC, cDC2 subset expanded most, whereas the proportion of cDC1 subset was reduced (Suppl. Fig. 6D-E). This shift in DC composition correlated with a trend toward enhanced activation of CD4⁺ T cells and an increase in Treg population in the mLN in CAC, suggesting that the altered DC landscape may promote a tolerogenic or regulatory environment rather than a strong cytotoxic CD8⁺ T cell response.

CD8⁺ T cells exhibit a mild increase in granzyme levels throughout CAC progression, peaking at day 60 (Suppl. Fig. 6F-G). This temporal pattern suggests a gradual enhancement of their cytotoxic potential as the disease advances toward malignancy, possibly reflecting an adaptive response to the evolving tumour microenvironment. Notably, although GZMB is among the most abundant proteins in CD8^+^ T cells, its concentration remained unchanged in mLN during CAC, similar to perforin (PFR1) levels (Suppl. Fig. 6G, 6H). This observation aligns with previous findings showing that GZMB is highly translated in both proliferating and non-proliferating T cells *in vivo*^24^.

All these findings suggest that cytotoxic activity in CD8⁺ T cells, together with immunosuppressive mechanisms mediated by Tregs, may be further reinforced in the tumour microenvironment, potentially contributing to immune evasion and cancer progression.

### Differential protein expression in the transition from colitis to cancer

Additional analysis to identify proteins with altered expression during the progression from colitis to cancer once again revealed more pronounced changes in CD4^+^ than in CD8^+^ T cells (Fig. 7A, 7B). Upregulated proteins in CD4^+^ T cells were mainly involved in biological processes related to the regulation of synaptic vesicle exocytosis and methylation, as well as ATP synthesis, as we described before (Fig. 5F, Fig. 7C).

**Figure 7.**
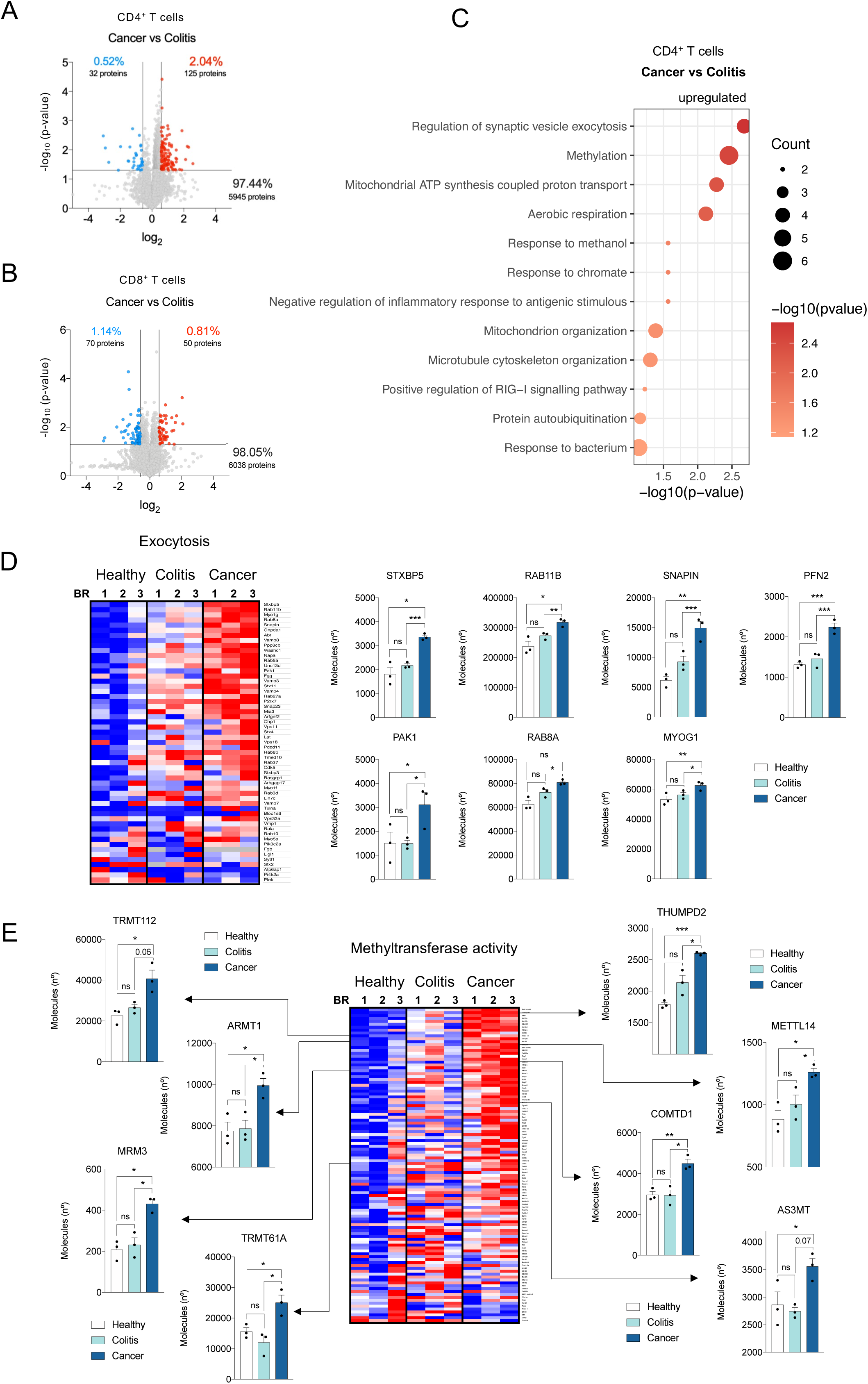
Proteomic differences between colitis and cancer in CD4⁺ and CD8⁺ T cells. Mice were treated with AOM/DSS, and at days 0 (Healthy), 15 (Colitis) and 60 (Cancer), CD4^+^ and CD8^+^ T cells were sorted from mLN. **(A-B)** Volcano plots showing the difference in the expression of proteins between Cancer and Colitis in (A) CD4^+^ T cells and (B) CD8^+^ T cells. The horizontal lines indicate a *p*-value of 0.05. The vertical lines indicate a fold change of ± 1.5. The percentage and number of proteins that are downregulated (blue), upregulated (red), or unchanged (grey) are indicated. **(C)** Bubble plot showing the enrichment analysis of GO biological processes of the differentially upregulated proteins between Cancer and Colitis in CD4^+^ T cells. Bubble size indicates the number of genes involved in each process, and bubble colour reflects the fold change between Cancer and Colitis. **(D)** Left: Heat map of proteins involved in the exocytosis process (GO:0006887) identified in CD4^+^ T cells proteomes during CAC progression. BR, biological replicate. Relative protein abundance is graded from low (blue) to high (red) to allow comparisons. Right: Copy numbers per cell (Molecules (n°)) of the proteins associated with the exocytosis process STXBP5, RAB11B, SNAPIN, PFN2, PAK1, RAB8A and MYOG1 in CD4^+^ T cells. Each dot represents a pool of three mice (n=3 pools). **(E)** Heat map of proteins and protein copy numbers involved in Methyltransferase activity (GO:0008168) identified in CD4^+^ T cells in AOM/DSS-treated mice. Relative protein abundance is graded from low (blue) to high (red) per row. BR, biological replicate. Protein copy numbers per cell (Molecules (n°)) of a selection of molecules are shown. Each dot represents a pool of three mice (n=3 pools). All the data are shown as mean ± SEM. In all panels, ns = non-significant, \**p* ≤ 0.05, \*\**p* ≤ 0.01, \*\*\**p* ≤ 0.001.

Increased levels of proteins like RAB11B, RAB8A STXBP5, SNAPIN or PAK1 (Fig. 7D), which are implicated in exocytosis, in cancer compared to colitis, may reflect an enhanced capacity of these cells to secrete cytokines and other effector molecules, thereby amplifying local inflammation and supporting/promoting a pro-tumorigenic microenvironment.

Methylation-related proteins can influence CD4^+^ T cells activation and differentiation by modulating diverse transcriptional programs. We found that proteins of methyltransferase complexes and methyltransferase cofactors such as METTL14, TRMT112, TRMT61A, ARMT1, MRM3, THUMPD2, AS3MT or COMTD1 were consistently upregulated in CD4^+^ T cells in cancer compared to colitis or healthy conditions (Fig. 7E). These findings suggest that CD4^+^ T cells undergo epigenetic reprogramming associated with the transition from inflammation to tumorigenesis. In cancer, altered methylation pathways may impair effector T cell function, promoting immune suppression and tumour progression^25^.

## Discussion

This study provides a comprehensive proteomic analysis of CD4⁺ and CD8⁺ T cells during the progression of colitis-associated colorectal cancer (CAC), profiling over 6000 proteins. Our data reveal strikingly different activation dynamics and metabolic adaptation between these T cell populations, highlighting the immunometabolic plasticity of CD4⁺ T cells and the restrained activation of CD8⁺ T cells. By analysing CD4⁺ and CD8⁺ T cells from mLNs, key sites for immune cell priming and activation in the gut, we captured early stages of T cell activation and differentiation before their migration into the tumour.

Upon treatment, CD4⁺ T cells, but not CD8⁺ T cells, showed increased protein synthesis, resulting in greater cell size and expansion. This was accompanied by higher expression of ribosomal proteins, amino acid transporters, and initiation factors, consistent with enhanced translational. These findings extend previous *in vitro* observations^26,27^ to an *in vivo* setting, demonstrating that chronic inflammation drives similar anabolic programs during CAC progression.

CD4⁺ T cell activation promotes clonal expansion and proliferation^28,29^, supported by the upregulation of DNA replication machinery and cell cycle regulators. These processes were linked to metabolic rewiring in CD4⁺ T cells, with enhanced mitochondrial mass, glycolysis, and expression of proteins implicated in nutrient uptake and ATP synthesis. Together, these adaptations provide the energy and biosynthetic resources required for sustained T cell growth during CAC. However, during the colitis-to-cancer transition, CD4⁺ T cells show a non-significant reduction in aerobic capacity, suggesting a metabolic shift away from mitochondrial oxidative phosphorylation. This progressive loss of metabolic plasticity may contribute to immune dysfunction and tumour immune evasion during CAC progression, thereby supporting the progression of the disease, as previously described in tumour-infiltrating lymphocytes^30,31^.

Beyond metabolic remodelling, our data point to additional layers of regulation in CD4⁺ T cells, including epigenetic and epitranscriptomic mechanisms that may sustain their proliferative and immunosuppressive programs during CAC progression. Several methylation-related proteins appear to play dual roles, contributing not only to DNA and RNA methylation but also to ribosomal RNA (rRNA) methylation, potentially enhancing ribosome biogenesis and global protein translation, a hallmark of highly proliferative cells within the tumour microenvironment. Notably, METTL14, a core component of the m⁶A methylation machinery, has been implicated in T cell activation and Treg function^32,33^. This observation aligns with our findings of increased Treg population and elevated FoxP3, ICOS, and CTLA-4 expression, suggesting that METTL14-driven epitranscriptomic modifications may reinforce immunosuppressive mechanisms and facilitate immune evasion during CAC progression.

Interestingly, despite both CD4⁺ and CD8⁺ T cells were exposed to similar inflammatory and oncogenic cues, CD8⁺ T cells maintained a more stable proteome, with minimal changes in protein content and in activation-related pathways. Nevertheless, during CAC, these cells showed reduced expression of TCA cycle components, indicating metabolic impairment and suggesting limited cytotoxic potential. This aligns with previous studies reporting dysfunctional CD8⁺ T cells in CAC and other inflammation-driven cancers^34^.

Proteomic profiles further indicate that CD4⁺ T cells differentiate into inflammatory Th1 and Treg subsets during colitis, persisting in cancer. In contrast, CD8⁺ T cells retain their cytotoxic identity, evidenced by elevated granzyme expression. Notably, despite minimal metabolic changes, granzyme production does not require enhanced metabolic activity, as previously described^35^, yet the functional activity of these CD8⁺ T cells in cancer appears limited. One possible mechanism involves the altered composition of DC subsets in the mLN, which influence CD4⁺ and CD8⁺ T cell activation and function. Moreover, in CD4^+^ T cells, increased cytokine production and upregulation of MAPK/NFκB pathways, key modulators of immune responses and T cell effector functions under inflammatory conditions, support their role in sustaining inflammation and promoting tumour progression, consistent with their known contribution to shaping the inflammatory microenvironment in CAC^36^.

Our data also reveal dynamic changes in accessory functions: both CD4⁺ and CD8⁺ T cells expressed MHC class II early in disease, but this declined during CAC. Given that activated T cells, particularly CD4⁺ subsets, can upregulate MHCII and act as antigen-presenting cells under inflammatory conditions^37–39^, amplifying immune responses through antigen-specific interactions, the loss of MHCII in CAC would suggest a shift toward immunosuppression, reducing antigen presentation and favouring tumour immune evasion.

Collectively, our findings provide new insights into the dynamic proteomic remodelling of T cells in mLN and emphasise the importance of CD4⁺ T cell metabolism and activation in Inflammatory bowel disease-associated cancer. Interventions aimed at restoring CD8⁺ T cell activity or controlling CD4⁺ T cell-driven inflammation appear to be promising therapeutic approaches. Specifically, enhancing mitochondrial function or glycolytic flexibility in CD8⁺ T cells, while limiting excessive activation in CD4⁺ T cells, warrants further investigation. Future studies should validate these observations in human samples and assess whether metabolic signatures can serve as biomarkers for CAC progression or therapeutic response. Modulating CD4⁺ T cell activity during early colitis could help prevent CAC development, whereas boosting CD8⁺ T cell responses may improve anti-tumour immunity in established solid tumours with an immunologically “cold” phenotype, such as CRC.

### Conclusion

Our findings reveal distinct immune adaptations of CD4⁺ and CD8⁺ T cells during CAC progression. Chronic inflammation drives CD4⁺ T cell activation and metabolic reprogramming, marked by increased glycolysis, mitochondrial biogenesis, and translational capacity, supporting sustained cytokine production and proliferation. In contrast, CD8⁺ T cells fail to undergo similar adaptation, showing impaired oxidative phosphorylation and reduced cytotoxic potential. This imbalance may create an immune environment that favours tumour growth while limiting effective antitumor responses. The dynamic interplay between these subsets underscores the need for targeted strategies to restrain CD4⁺ T cell-driven inflammation early and restore CD8⁺ T cell fitness in established tumours.

## Material and Methods

### Mice

C57BL/6J wild-type mice were used in this study. Mice were housed in specific-pathogen-free conditions in accordance with European Union regulations; work was performed according to the regulation of local CNB and CSIC bioethics committees and approved by the Community of Madrid (PROEX 071/19 and 289.2/20).

### Animal models and *in vivo* procedures

Colitis-associated colorectal cancer (CAC) was induced as described^13^. Male and female 10-14 weeks old mice were intraperitoneally injected with AOM (10 mg/kg; Sigma-Aldrich, A5486). On day 6 after AOM injection, mice underwent a 5-day treatment with 2% DSS (w/v) (MW 36,000-50,000 Da; MP Biomedicals, 160110) in drinking water, followed by 15 days administration of normal drinking water. This treatment was repeated twice. Mice were sacrificed at the indicated times (0 days, healthy conditions, 15 days, colitis, and 60 days, colitis-associated colon cancer) mesenteric lymph nodes (mLN) and colon were removed for analysis. In all procedures, mice were monitored for signs of distress, weight loss and rectal bleeding. For the histological images the colon were formalin-fixed, paraffin-embedded and stained for haematoxylin and eosin.

### T cell isolation

To analyse T cell proteome, for *ex vivo* naïve CD4^+^ and CD8^+^ T cells, mLN from three mice were pooled and used per biological replicate (BR). Three BR were generated per condition. Single cell suspensions were generated by mashing mLNs in RPMI medium and filtered through a 70 μm cell strainer. Cells were harvested by centrifugation and prepared for cell sorting to isolate pure CD4^+^ and CD8^+^ T cell populations.

To analyse cytokine production, CD4^+^ and CD8^+^ T cells purified from mLNs by negative selection (MojoSort Mouse CD4 T Cell Isolation Kit, BioLegend 480033, or MojoSort Mouse CD8 T Cell Isolation Kit, BioLegend, 480035), harvested by centrifugation and prepared for RNA extraction to quantify cytokine mRNA expression by qPCR.

### Gene expression analysis

For mRNA analysis, isolated CD4^+^ or CD8^+^ T cells from mLN in healthy, colitis and cancer conditions were lysed with NZYol (NZYtech, MB18501) and RNA extracted using a standard protocol with chloroform-isopropanol-ethanol. cDNA for real-time quantitative PCR (qPCR) was generated from total RNA using the High Capacity cDNA Reverse Transcription Kit (Applied Biosystems, 4368814) and as described^40,41^. PCR reactions were carried out in a QuantStudio5 (Applied Biosystems) and SDS v2.2 software was used to analyse the results by the Comparative Ct Method (ΔΔCt). X-fold change in mRNA expression was quantified relative to the control *Actb* mRNA. The different genes analysed in this work were amplified using the primers described in (Suppl. Table 1).

### Analysis of Cell Bioenergetics

The oxygen consumption rate (OCR) and the extracellular acidification rate (ECAR) were measured using an XFe96 Extracellular Flux analyser (Seahorse Bioscience). CD4^+^ or CD8^+^ T cells were seeded (0.4 x 10^6^ cells/well) on a XFe96 cell culture microplate previously treated with Cell-Tak (Corning). Mitochondrial stress tests were performed by incubating cells for 1 h in the absence of CO_2_ in non-buffered XFe assay medium pH 7.4 (Seahorse Bioscience), supplemented with 25 mM glucose, 2 mM glutamine and 1 mM sodium pyruvate. MitoStress test was conducted following previously described protocols, with minor modifications. Briefly, after basal rate measurements, different modulators of mitochondrial respiration were injected sequentially (1): 2.5 µM oligomycin to inhibit ATP-synthase and to calculate the ATP-linked oxygen consumption (2); 1.3 µM carbonyl cyanide-P-trifluoro-methoxyphenylhydrazone (FCCP; an uncoupling agent) to obtain the maximum respiration under stress conditions; and (3) a mix of 0.5 µM rotenone/antimycin A to completely block mitochondrial respiration by inhibiting complexes I and III respectively. Glycolysis stress tests were performed using cells starved in a non-CO_2_ incubator for 1 h at 37°C in non-buffered XF assay medium (Seahorse Bioscience) supplemented with 2 mM glutamine and 1 mM sodium pyruvate. After measuring basal ECAR, 15 mM glucose were injected to stimulate glycolysis, followed by 2.5 µM oligomycin to obtain the maximum glycolytic capacity after the inhibition of oxygen consumption. Finally, 100 mM of 2-deoxy-D-glucose (2-DG) were injected to shut down glycolysis. OCR and ECAR were measured three times after the addition of each drug. At least four animals per condition were used in each experiment. Calculations were performed with the Seahorse XFe Cell Test Report Generator software (Seahorse Bioscience).

### Flow cytometry analysis and cell sorting

Flow cytometry analysis was performed with 1 x 10^6^ cells per condition resuspended in 3% (v/v) FBS-PBS; cells were stained with combinations of fluorescently labelled antibodies to the cell surface and to intracellular or intranuclear markers, indicated in the figures and in (Suppl. Table 2), and incubated at 4°C for 30 min. For intranuclear staining, cells were incubated with a commercial kit (Foxp3/Transcription Factor Staining buffer Set, eBioscience, 00-5523-00) according to the manufacturer’s instructions. The analysis of cell populations was performed in a in Gallios Analyzer (Beckman Coulter) using the software FlowJo v10 (FlowJo). Cell sorting was performed on a FACSAria Fusion (BD Biosciences). Staining was carried out in PBS supplemented with 1% (v/v) FBS. Cells were stained with fluorophore-conjugated antibodies anti-CD4, -CD8, -Thy1.2(CD90.2) and DAPI. Ex vivo CD4^+^ and CD8^+^ T cells were sorted from mLNs of untreated and AOM/DSS-treated mice as DAPI^−^Thy1.2^+^CD4^+^CD8^−^ or DAPI^−^Thy1.2^+^CD4^−^CD8^+^. Three mice were used per BR. All sorted cells were then washed twice with HBSS before being snap frozen in liquid nitrogen and stored at -80°C until further processing. All cell sorting was performed in 1% (v/v) FBS-PBS and cells were collected in RPMI1640 1% (v/v) FBS.

### Proteomics sample preparation

After sorting, T cell pellets were processed using S-trap micro protocol (Protifi) following manufacturer instructions with little modification. After application of the samples on the S-trap micro spin column, trapped proteins were washed three times with S-trap binding buffer. Digestion with trypsin was carried out overnight at 37°C in 150 μl of ammonium bicarbonate buffer (TEAB) at 50 mM. Elution of peptides from S-trap micro spin column was achieved by centrifugation (4000 x g, 1 min) by adding 40 μl of 50 mM ammonium bicarbonate, then 40 μl of 0.2% (v/v) aqueous formic acid and finally 40 μl of 50% (v/v) acetonitrile/0.2% (v/v) formic acid. Resulting tryptic peptides were dried using a Speed-vac machine overnight and quantified by CBQCA method (ThermoFisher, C667) following manufacturer instructions.

### Proteomics sample acquisition for single-shot DIA analysis

Peptides were diluted with 1% (v/v) formic acid to a final concentration of 0.1 μg/μl for mass spectrometry analysis and were submitted to the FingerPrints Proteomic Facility at the University of Dundee. Samples were run on an Ultimate 3000 RSL Cnano system (Thermo Scientific) coupled to an Exploris 480 Mass Spectrometer (Thermo Scientific). Samples run in Ultimate 3000 recommended buffer conditions comprising: Buffer A: 0.1% (v/v) formic acid and Buffer B: 80% (v/v) acetonitrile in 0.1% (v/v) formic acid. Samples were injected (15 μl/min) and washed onto a C18 trap with 0.1% (v/v) formic acid (Buffer C). After 5 min a gradient formed with buffers A and B. Peptides were initially trapped on an Acclaim PepMap 100 (C18, 100 μM x 2 cm) and then separated on an Easy-Spray PepMap RSLC C18 column (75 μM x 50 cm) (Thermo Scientific). The samples were then transferred to the mass spectrometer via an Easy-Spray source with the temperature set at 50 .C and a source voltage of 1.9 kV. A 140 min acquisition was run on the mass spectrometer which was operating in data-independent acquisition mode with 1 MS1 scan followed by 45 DIA scans. Peptide application mode was selected and run for a total of 140 min.

### Analysis and processing of proteomic data

Quantification of reporter ions was carried out using Spectronaut version 16 (Biogosis) using a direct DIA analysis with Trypsin set as the digestive enzyme. Protein N-terminal acetylation, methionine oxidation and asparagine deamination were selected as variable modification, with carbamidomethylation of cysteine residues selected as a fixed modification. A false discovery rate of 1% was set for identification. Copy numbers for identified proteins were estimated using the histone ruler method originally published by using Perseus software supplied by MaxQuant (version 1.6.10.43). Copy numbers of histones in a diploid mouse cell get assigned to the summed peptide intensities of all histones present in a sample. The ratio between the histone peptide intensity and the summed peptide intensities of the other identified proteins is then used to estimate copy number per cell for all identified proteins in the data set. The mass contribution of proteins (pg/cell) was calculated as (protein copy number) x (molecular weight (Daltons)) / (Avogadro’s constant).

### Statistical analysis

For the *in vivo* analysis, group sizes were established according to the Spanish ethical legislation. For the proteomic analysis, *p*-values were calculated using two-tailed *t*-test with unequal variance on log_10_ transformed copy number per cell values, using Microsoft Excel. Volcano plots were generated by plotting negative log_10_-transformed *p*-values against log-transformed fold changes. Proteins with a *p*-value < 0.05 and a fold change in expression greater than 1.5 between conditions were considered significantly differentially expressed. Pathway analysis of these proteins was performed using the online tool DAVID (https://david.ncifcrf.gov/summary.jsp). Additional data analyses were conducted using Student’s *t*-test in GraphPad Prism software. In all cases, *p*-values < 0.05 were considered statistically significant. Data are shown as mean ± SEM. For clarity, non-significant differences are not shown in some figures. Images and schemes were created using Biorender.com.

**Table 1.**
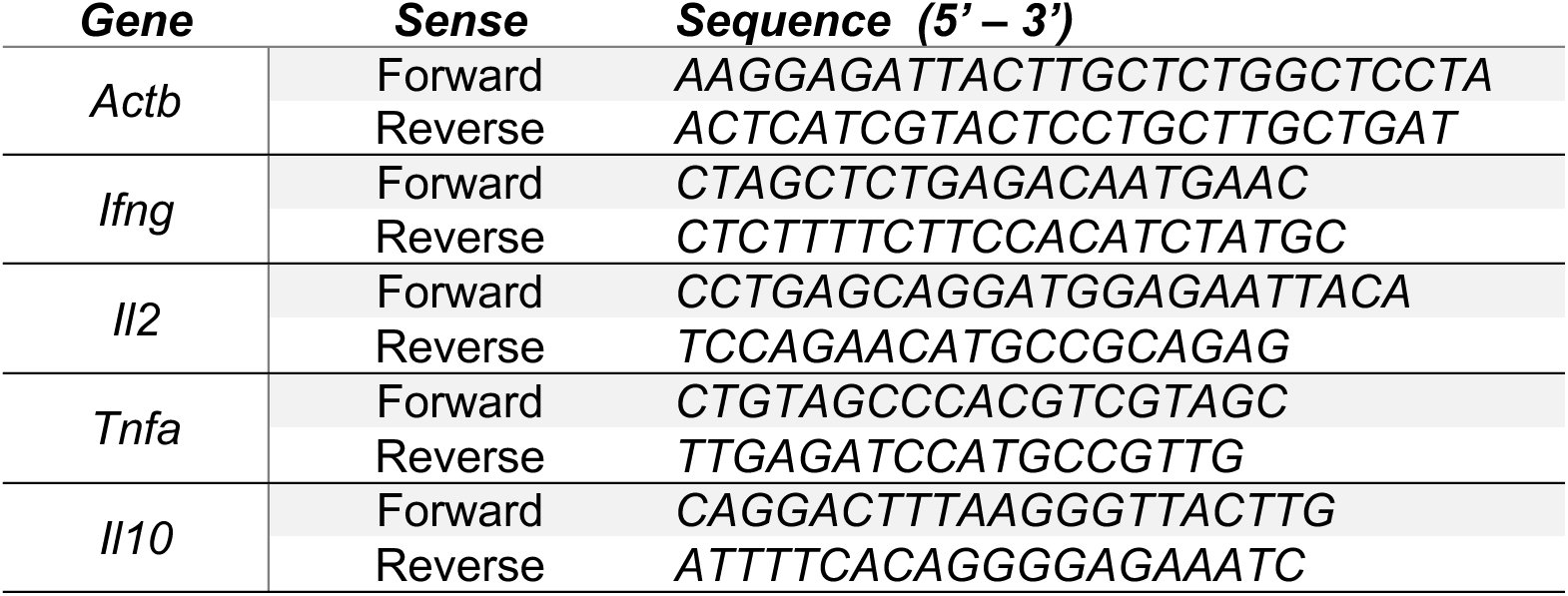
Primers used for RT-qPCR analysis.

**Table 2.** Antibodies used for Flow cytometry analysis.

| **Antibody** | **Source** | **Identifier** | **Concentration** | **Dilution** |
| --- | --- | --- | --- | --- |
| *CD90.2-FITC* | eBioscience | 11-0902-85 | 0.125 mg/ml | 1/100 |
| *CD4-APCe780* | eBioscience | 47-0042-82 | 0.2 mg/ml | 1/100 |
| *CD8-PERCP* | Biolegend | 100732 | 0.05 mg/ml | 1/100 |
| *CD45-PERCPCy5.5* | Biolegend | 103132 | 0.05 mg/ml | 1/100 |
| *FOXP3-ef450* | eBioscience | 48-5773-80 | 0.2 mg/ml | 1/50 |
| *CD19-AlexaFluor700* | eBioscience | 56-0193-82 | 0.05 mg/ml | 1/100 |
| *CD45-BV570* | Biolegend | 103136 | 0.2 mg/ml | 1/100 |
| *CD4-AlexaFluor700* | eBioscience | 56-0042-82 | 0.025 mg/ml | 1/100 |
| *CD8-ef450* | eBioscience | 48-0081-82 | 0.05 mg/ml | 1/100 |
| *IA/IE-FITC* | BioLegend | 107606 | 0.125 mg/ml | 1/100 |
| *Ly6C-PECF594* | BD Biosciences | 562728 | 0.2 mg/ml | 1/100 |
| *F4/80-PECy5* | eBioscience | 15-4801-82 | 0.05 mg/ml | 1/100 |
| *CD11c-PECy7* | eBioscience | 25-0114-81 | 0.05 mg/ml | 1/100 |
| *CD11b-APC* | BioLegend | 101212 | 0.2 mg/ml | 1/100 |
| *CD45-AlexaFluor700* | BioLegend | 103127 | 0.125 mg/ml | 1/100 |
| *Ly6G-APCe780* | eBioscience | 47-9668-82 | 0.2 mg/ml | 1/100 |
| *Live/Dead Fixable Violet* | Invitrogen | L34963 |  |  |
| *Live/Dead Fixable Red* | Invitrogen | L34972 |  |  |

## Acknowledgments

We thank R. Gómez-Caro for technical support, and J. Isoler-Alcaraz for assistance with figure preparation using R. We also thank the Flow Cytometry, Animal, and Histology Facilities at CNB-CSIC, the Flow Cytometry Facility at CBMSO, and the FingerPrints Proteomics Facility at the University of Dundee for technical assistance. This research was funded by MICIU/AEI/10.13039/501100011033 (PID2022-139944OB-I00) to AC and JJSE, Wellcome Trust Principal Research Fellowship grant to DAC (205023/Z/16/Z), and (PID2022-138295OB-I00) to MGC. PF and DG-R received MCIN FPI fellowships, and ED-M received a MEFP FPU fellowship. DG-R also received an EMBO Scientific Exchange Grant (9406). We have used Co-pilot to reduce the extension of the manuscript and to fit within the formatting requirements.

## Contributions

DG-R and LS performed proteomic and bioinformatic analyses. DG-R, PF and ED-M contributed to immunohistochemistry and flow cytometry analyses. DG-R and MGdC performed seahorse analyses. JJS-E, MGdC, DAC, and AC provided essential resources. DG-R, LS, DAC, and AC conceived and supervised the study. PF drafted figure legends, and DG-R and AC wrote the manuscript with input from all authors. All authors reviewed and approved the final version of the manuscript.

## Ethics approval

All animal experiments were carried out at the facilities of the National Centre for Biotechnology (CNB-CSIC) in accordance with national and European guidelines. Work was performed according to the regulation of local CNB and CSIC bioethics committees and approved by the Community of Madrid PROEX 071/19 and 289.2/20.

## Conflicts of interest

The authors declare that they have no conflict of interest.

## Notes

### Competing Interest Statement

The authors have declared no competing interest.

### Summary of Updates

Typo mistakes and supplementary Fig 1

