## Supplementary Information for "Differential proteomic and metabolic remodelling of CD4⁺ and CD8⁺ T cells in mesenteric lymph nodes during colitis-associated colorectal cancer"

**Supplementary Figure 1. Analysis of immune populations during CAC progression**

**(A)** Schematic representation of CAC model and the protocol followed to obtain sorted $CD4^+$ and $CD8^+$ T cells from mLN of healthy, colitic, and cancer-bearing mice. **(B)** Mice were treated with AOM/DSS for 0, 15 or 60 days. Representative images of formalin-fixed, paraffin-embedded Healthy, Colitis and Cancer colon tissues stained with hematoxylin and eosin. Ulcers, infiltration area and tumours are outlined with dashed yellow lines. **(C)** Mice were treated as in (B). mLN cells were stained with antibodies and analysed by flow cytometry. Representative flow cytometry plots showing the gate strategy used. **(D)** Total cell number of Dendritic cells ($CD45^+$ $CD11c^+$), Monocytes ($CD45^+$ $CD11b^+$ $Ly6C^+$), Neutrophils ($CD45^+$ $CD11b^+$ $Ly6G^+$), Macrophages ($CD45^+$ $CD11b^+$ $F4/80^+$) and B cells ($CD45^+$ $CD19^+$) in mLN from healthy and cancer mice. Each dot represents a single mouse (n=3-5). All the data are shown as mean $\pm$ SEM. In all panels, ns = non-significant, $*p \leq 0.05$.

**Supplementary Figure 2. Analysis of biological processes differentially expressed in $CD4^+$ and $CD8^+$ T cells in Colitis and Healthy stage**

Mice were treated with AOM/DSS, and at days 0 (Healthy), 15 (Colitis) and 60 (Cancer), $CD4^+$ and $CD8^+$ T cells were sorted from mLN. **(A-B)** Bubble plots showing the enrichment analysis of GO biological processes of the differentially upregulated proteins and downregulated proteins in Colitis versus Healthy in (A) $CD4^+$ and (B) $CD8^+$ T cells. Bubble size indicates the number of genes involved in each process, and bubble colour reflects the fold change in Colitis versus Healthy conditions.

**Supplementary Figure 3. Analysis of molecules involved in antigen presentation**

Mice were treated with AOM/DSS, and at days 0 (Healthy), 15 (Colitis) and 60 (Cancer), $CD4^+$ and $CD8^+$ T cells were sorted from mLN. **(A-B)** Copy numbers per cell (Molecules (nº)) of the molecules involved in antigen processing and presentation via MHC-II in (A) $CD4^+$ T cells and (B) $CD8^+$ T cells. **(C, E)** Representative flow cytometry plots (Left panels). Percentage of MHCII$^+$ CD4$^+$ T cells (C), percentage of MHCII$^+$ CD8$^+$ T cells (E) in mLN from healthy and colitic mice (Right panels). **(D, F)** Representative flow cytometry histograms (Left panels). Levels of MHCII expression (MFI) in CD4$^+$ T cells (D) and in CD8$^+$ T cells (F) (Right panels). Each dot represents a pool of three mice (n=3 pools). All the data are shown as mean ± SEM. In all panels, ns = non-significant, *$p \leq 0.05$, **$p \leq 0.01$, ***$p \leq 0.001$.

**Supplementary Figure 4. Analysis of proteins involved in DNA replication and cell-cycle during CAC**

Mice were treated with AOM/DSS, and at days 0 (Healthy), 15 (Colitis) and 60 (Cancer), CD8$^+$ T cells were sorted from mLN. Protein copy numbers per cell (Molecules (nº)) of a selection of molecules related with DNA replication and cell cycle are shown. Each dot represents a pool of three mice (n=3 pools). All the data are shown as mean ± SEM. In all panels, ns = non-significant, *$p \leq 0.05$, **$p \leq 0.01$.

**Supplementary Figure 5. Analysis of glucose metabolism of CD8$^+$ T cells during CAC**

Mice were treated with AOM/DSS, and at days 0 (Healthy), 15 (Colitis) and 60 (Cancer), CD8$^+$ T cells were sorted from mLN. **(A)** Total glycolytic enzymes mass per cell in CD8$^+$ T cells. Each dot represents a pool of three mice (n=3 pools). **(B)** Expression profiles of glycolytic enzymes during CAC progression. Data are represented as protein copy numbers per cell (Molecules (nº)) in CD8$^+$ T cells. Each dot represents a pool of three mice (n=3 pools). **(C-D)** Abundances of (C) the glucose transporters SLC2A1 and SLC2A3 and (D) the lactate transporter SLC16A1 in CD8$^+$ T cells. Data are represented as protein copy numbers per cell (Molecules (nº)). Each dot represents a pool of three mice (n=3 pools). **(G)** Real-time monitorization of extracellular acidification rate (ECAR) in CD8$^+$ T cells along CAC progression (n=5 mice per condition). Arrow indicates the time of injection of glucose. **(H)** ECAR profile of CD8$^+$ T cells in basal state in healthy, colitis, and CAC (n= 5). **(I)** ECAR profile of CD8$^+$ T cells before (-) and after (+) glucose treatment (n= 5). All the data are shown as mean ± SEM. In all panels, ns = non-significant, *$p \leq 0.05$.

**Supplementary Figure 6. Proteomic and immunophenotypic analysis of CD8$^+$ T cells and dendritic cells in CAC**

**(A)** Volcano plot showing the differential expression of proteins associated with T cell activation (GO:0042110) and DNA-binding transcription factor activity (GO:0003700) in cancer versus healthy CD8$^+$ T cells. The horizontal lines indicate a *p*-value of 0.05. The vertical lines indicate a fold change of ± 1.5. **(B)** Mice were treated with AOM/DSS for 0, 15 or 60 days. mLN cells were stained with antibodies and analysed by flow cytometry. Representative flow cytometry plots. **(C-E)** Percentage of plasmacytoid dendritic cells (pDC), monocytic DC (moDC), DC, DC-cDC1, and DC-cDC2 populations in mLN from Healthy and Cancer mice are shown (n=3-7 mice/group). **(F)** Heat map of granzymes identified in CD8$^+$ T cells in AOM/DSS-treated mice. Relative protein abundance is graded from low (blue) to high (red) per row. BR, biological replicate. **(G, H)** Protein copy numbers per cell (Molecules (nº)) of a selection of molecules are shown. Each dot represents a pool of three mice (n=3 pools). All the data are shown as mean ± SEM. In all panels, ns = non-significant, *$p \leq 0.05$, **$p \leq 0.01$.

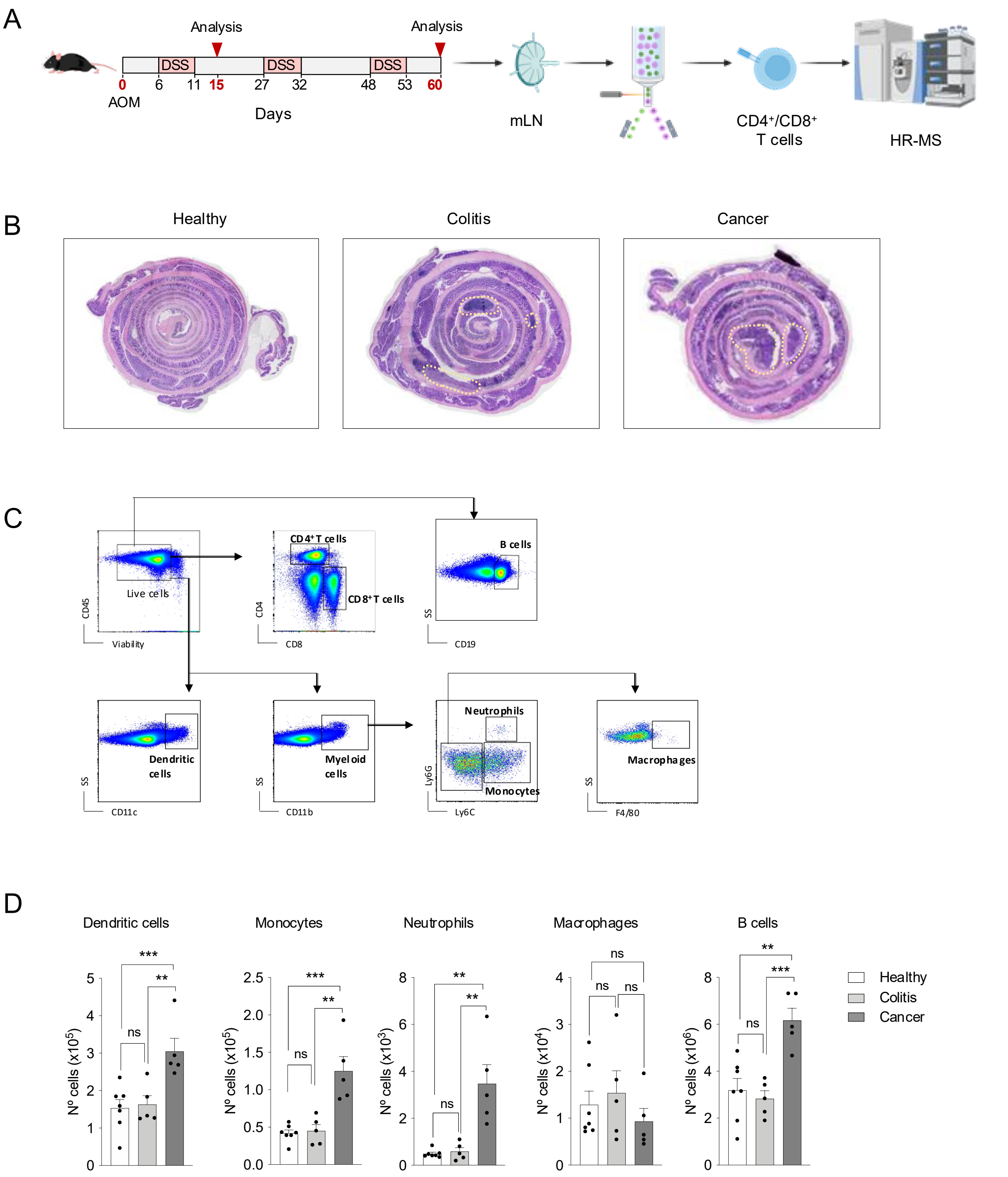
A
Analysis
Analysis
DSS
DSS
DSS
0
6
11
15
27
32
48
53
60
AOM
Days
mLN
CD4⁺/CD8⁺
T cells
HR-MS
B
Healthy
Colitis
Cancer
C
Live cells
CD45
Viability
CD4⁺T cells
CD8⁺T cells
CD4
CD8
B cells
SS
CD19
Dendritic cells
SS
CD11c
Myeloid cells
SS
CD11b
Neutrophils
Monocytes
Ly6G
Ly6C
Macrophages
SS
F4/80
D
Dendritic cells
Monocytes
Neutrophils
Macrophages
B cells
Nº cells (x10⁵)
Nº cells (x10⁵)
Nº cells (x10³)
Nº cells (x10⁴)
Nº cells (x10⁶)
Healthy
Colitis
Cancer

Suppl. Fig. 1

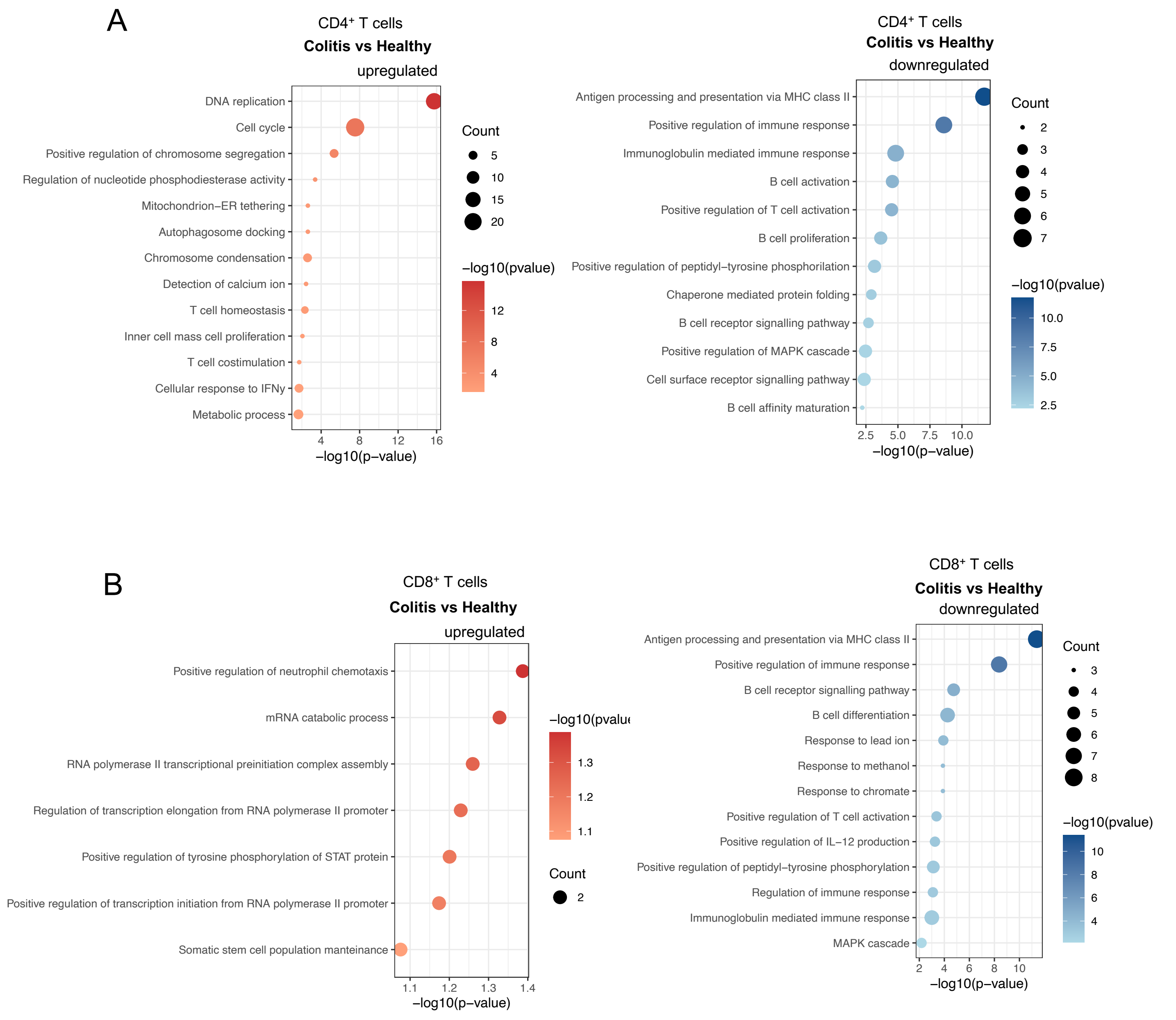

A
CD4⁺ T cells
Colitis vs Healthy
upregulated
DNA replication
Cell cycle
Positive regulation of chromosome segregation
Regulation of nucleotide phosphodiesterase activity
Mitochondrion–ER tethering
Autophagosome docking
Chromosome condensation
Detection of calcium ion
T cell homeostasis
Inner cell mass cell proliferation
T cell costimulation
Cellular response to IFNγ
Metabolic process
4 8 12 16
−log10(p−value)
Count
5
10
15
20
−log10(pvalue)
12
8
4
CD4⁺ T cells
Colitis vs Healthy
downregulated
Antigen processing and presentation via MHC class II
Positive regulation of immune response
Immunoglobulin mediated immune response
B cell activation
Positive regulation of T cell activation
B cell proliferation
Positive regulation of peptidyl–tyrosine phosphorilation
Chaperone mediated protein folding
B cell receptor signalling pathway
Positive regulation of MAPK cascade
Cell surface receptor signalling pathway
B cell affinity maturation
2.5 5.0 7.5 10.0
−log10(p−value)
Count
2
3
4
5
6
7
−log10(pvalue)
10.0
7.5
5.0
2.5
B
CD8⁺ T cells
Colitis vs Healthy
upregulated
Positive regulation of neutrophil chemotaxis
mRNA catabolic process
RNA polymerase II transcriptional preinitiation complex assembly
Regulation of transcription elongation from RNA polymerase II promoter
Positive regulation of tyrosine phosphorylation of STAT protein
Positive regulation of transcription initiation from RNA polymerase II promoter
Somatic stem cell population manteinance
1.1 1.2 1.3 1.4
−log10(p−value)
1.3
1.2
1.1
Count
2
CD8⁺ T cells
Colitis vs Healthy
downregulated
Antigen processing and presentation via MHC class II
Positive regulation of immune response
B cell receptor signalling pathway
B cell differentiation
Response to lead ion
Response to methanol
Response to chromate
Positive regulation of T cell activation
Positive regulation of IL−12 production
Positive regulation of peptidyl−tyrosine phosphorylation
Regulation of immune response
Immunoglobulin mediated immune response
MAPK cascade
2 4 6 8 10
−log10(p−value)
Count
3
4
5
6
7
8
−log10(pvalue)
10
8
6
4


Suppl. Fig. 2

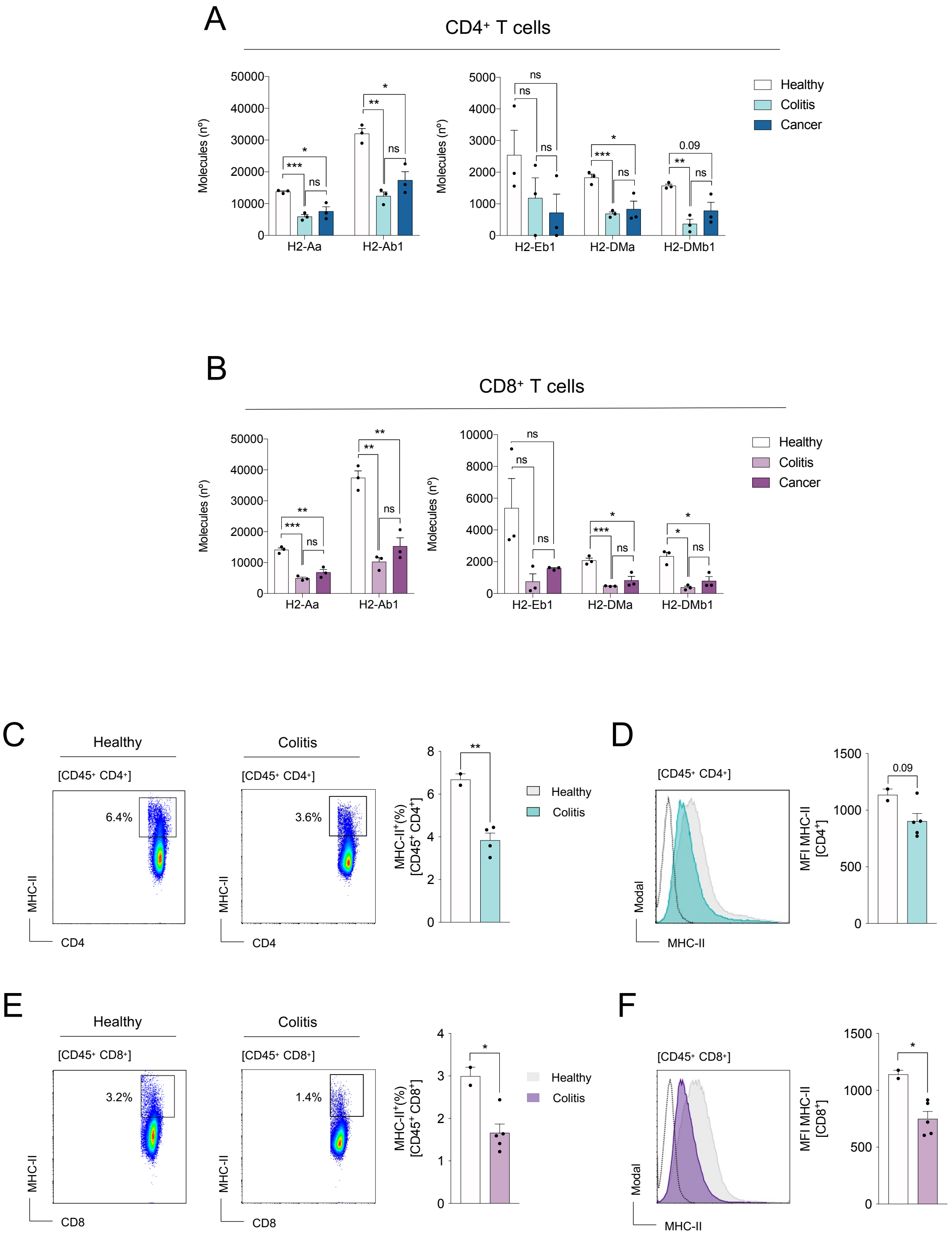

A
CD4⁺ T cells
Molecules (nº)
H2-Aa
H2-Ab1
H2-Eb1
H2-DMa
H2-DMb1
Healthy
Colitis
Cancer
B
CD8⁺ T cells
C
Healthy
Colitis
[CD45⁺ CD4⁺]
6.4%
3.6%
MHC-II
CD4
MHC-II⁺(%) [CD45⁺ CD4⁺]
D
Modal
MFI MHC-II [CD4⁺]
0.09
E
[CD45⁺ CD8⁺]
3.2%
1.4%
CD8
MHC-II⁺(%) [CD45⁺ CD8⁺]
F
MFI MHC-II [CD8⁺]


Suppl. Fig. 3

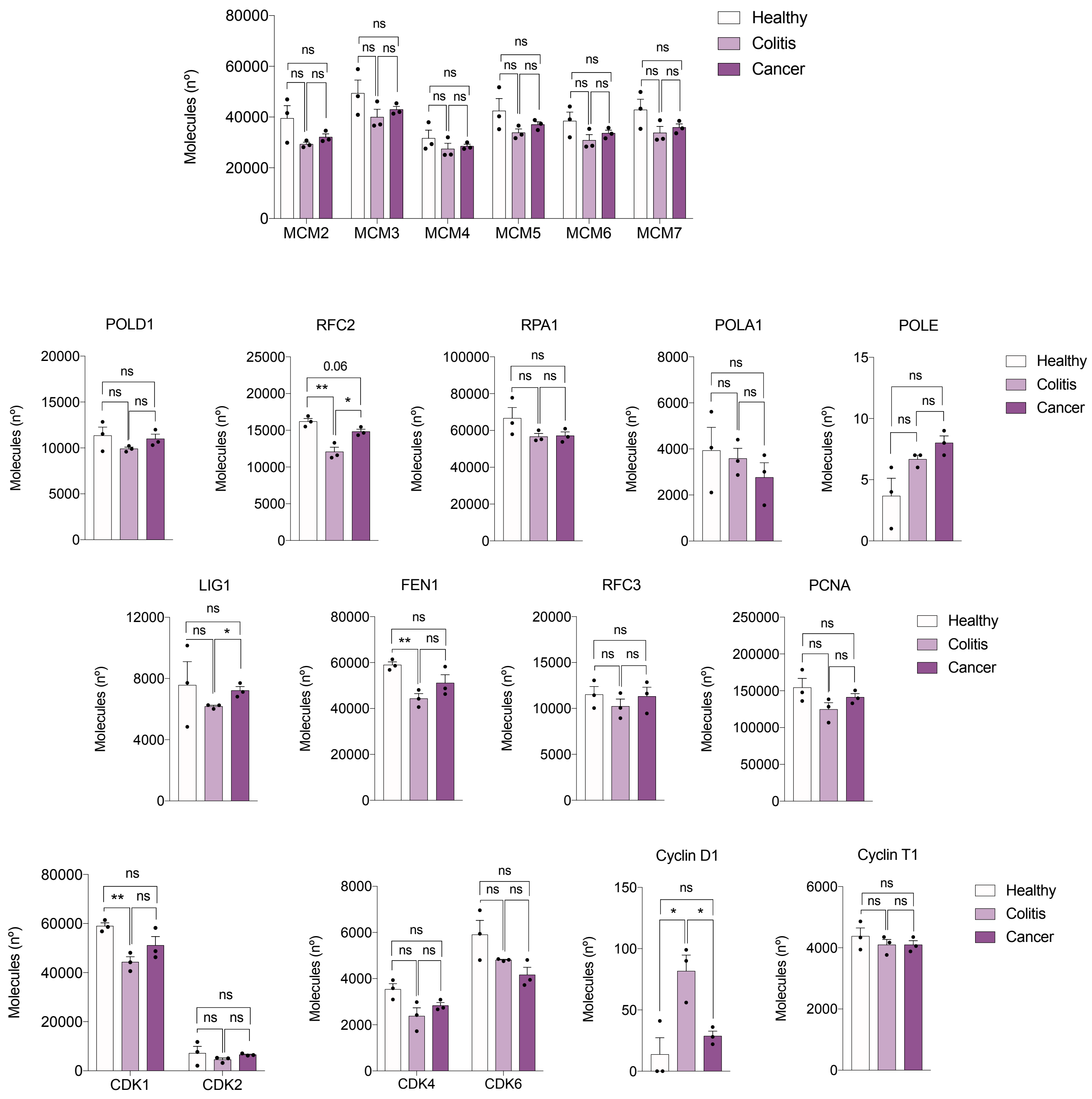
Molecules (nº)
MCM2
MCM3
MCM4
MCM5
MCM6
MCM7
Healthy
Colitis
Cancer
ns
POLD1
RFC2
0.06
**
*
RPA1
POLA1
POLE
LIG1
FEN1
RFC3
PCNA
CDK1
CDK2
CDK4
CDK6
Cyclin D1
Cyclin T1

Suppl. Fig. 4

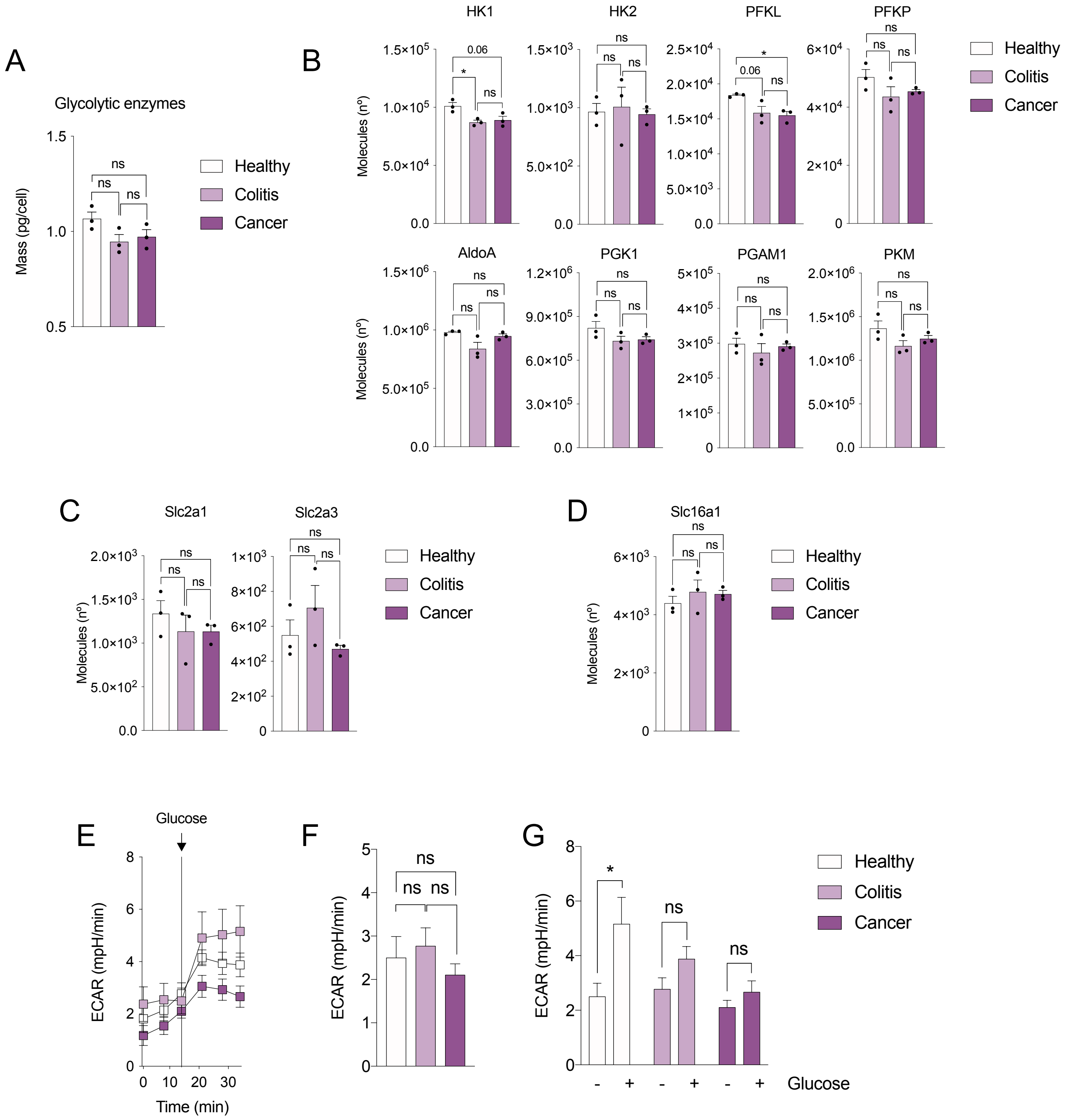
A
Glycolytic enzymes
Mass (pg/cell)
Healthy
Colitis
Cancer
B
HK1
HK2
PFKL
PFKP
AldoA
PGK1
PGAM1
PKM
Molecules (nº)
C
Slc2a1
Slc2a3
D
Slc16a1
E
Glucose
ECAR (mpH/min)
Time (min)
F
G
Glucose

Suppl. Fig. 5

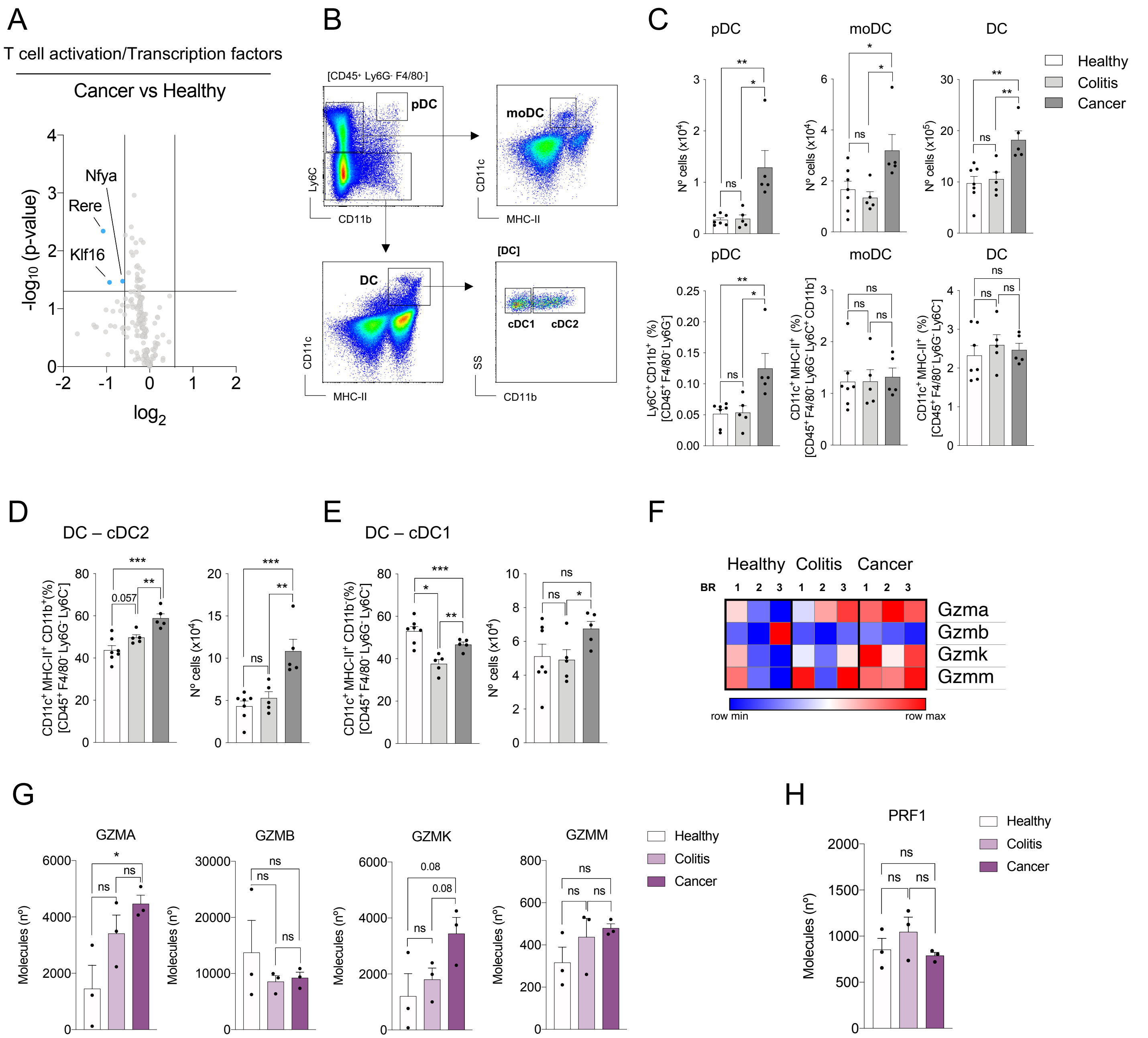
A
T cell activation/Transcription factors
Cancer vs Healthy
Nfya
Rere
Klf16
-log10 (p-value)
log2
B
[CD45+ Ly6G- F4/80-]
pDC
moDC
DC
[DC]
cDC1
cDC2
Ly6C
CD11b
CD11c
MHC-II
SS
C
pDC
moDC
DC
Healthy
Colitis
Cancer
Nº cells (x10^4)
Nº cells (x10^5)
Ly6C+ CD11b+ (%) [CD45+ F4/80- Ly6G-]
CD11c+ MHC-II+ (%) [CD45+ F4/80- Ly6G- Ly6C+ CD11b-]
CD11c+ MHC-II+ (%) [CD45+ F4/80- Ly6G- Ly6C-]
D
DC – cDC2
CD11c+ MHC-II+ CD11b+(%) [CD45+ F4/80- Ly6G- Ly6C-]
0.057
E
DC – cDC1
CD11c+ MHC-II+ CD11b-(%) [CD45+ F4/80- Ly6G- Ly6C-]
F
Healthy Colitis Cancer
BR
Gzma
Gzmb
Gzmk
Gzmm
row min
row max
G
GZMA
GZMB
GZMK
GZMM
Molecules (nº)
0.08
H
PRF1

Suppl. Fig. 6
